# The nucleoporin Nup50 activates the Ran guanyl-nucleotide exchange factor RCC1 to promote mitotic NPC assembly

**DOI:** 10.1101/2021.03.31.437874

**Authors:** Guillaume Holzer, Paola de Magistris, Cathrin Gramminger, Ruchika Sachdev, Adriana Magalska, Allana Schooley, Scheufen Anja, Birgitt Lennartz, Marianna Tatarek-Nossol, Hongqi Lue, Monika I. Linder, Ulrike Kutay, Christian Preisinger, Daniel Moreno-Andres, Wolfram Antonin

## Abstract

During mitotic exit, thousands of nuclear pore complexes (NPCs) assemble concomitant with the nuclear envelope to build a transport-competent nucleus. We show here that Nup50 plays a crucial role in NPC assembly that is independent of its well-established function in nuclear transport. RNAi-mediated downregulation in cells or immunodepletion of the protein in Xenopus egg extracts interferes with NPC assembly. We define a conserved central region of 46 residues in Nup50 that is crucial for Nup153 and MEL28/ELYS binding, and NPC interaction. Surprisingly, neither NPC interaction nor binding of Nup50 to importin α, β, the GTPase Ran or chromatin is crucial for its function in the assembly process. Instead, we discovered that an N-terminal fragment of Nup50 can stimulate the Ran guanine exchange factor RCC1 and NPC assembly, indicating that Nup50 acts via the Ran system in mitotic NPC reformation. In support of this conclusion, Nup50 mutants defective in RCC1 binding and stimulation cannot replace the wild type protein in in vitro NPC assembly assays.

## Introduction

Nuclear pore complexes (NPCs) are the gatekeepers of the nucleus, controlling the exchange of proteins and nucleic acids between cytoplasm and nucleoplasm. They are embedded in the nuclear envelope (NE) and with a mass of approximately 125 MDa the largest protein complexes in most vertebrate cells. Despite their enormous size they are only composed of about 30 different proteins called nucleoporins all present in multiple copies. Because of their common function in all nucleated cells, the general structure of NPCs is evolutionary conserved [1, 2]. Their core structure is formed by three rings embedded in the NE, a nucleoplasmic, an inner, and a cytoplasmic ring. This symmetric core encloses a central transport channel and anchors extensions on the cytoplasmic and nuclear faces forming cytoplasmic filaments and nuclear basket, respectively.

In metazoans, NPCs form at two different stages of the cell cycle by distinct mechanisms (for review see [3, 4]). During interphase, new NPCs assemble into the intact NE. This assembly pathway is thought to be initiated at the membranes of the NE, although the mechanistic details remain poorly defined. At the end of mitosis, NPC re-assembly occurs concomitantly with the formation of a closed NE and is initiated by the chromatin binding nucleoporin MEL28/ELYS directing mitotic NPC assembly to the surface of the decondensing chromatin [5, 6]. The small GTPase Ran serves as a second important spatial regulator that guides NPC assembly towards chromatin [7]. High RanGTP concentrations around chromatin, generated by the action of the chromatin-bound Ran guanine exchange factor (GEF) RCC1, are thought to release inhibitory nuclear transport factors from key nucleoporins which can then act in NPC assembly. The crucial functions of both MEL28/ELYS and the Ran system is probably best exemplified by experiments where tethering MEL28/ELYS and RCC1 to DNA beads is sufficient to initiate NPC assembly [8].

Cell free and cellular assays have revealed a picture of a step-wise assembly of the NPC core structure. Late in the assembly process, the cytoplasmic filaments and the nuclear basket structure form. The nuclear basket is comprised in vertebrates of three nucleoporins, Nup153, TPR, and Nup50 [9–11], also referred to as Npap60 [12], which have multiple roles in nuclear import and export of proteins, chromatin remodeling, control, of gene expression as well as RNA processing and export [13, 14]. Both Nup50 [10, 15, 16] and TPR [17] interact with Nup153. Accordingly, Nup50 NPC localization depends an Nup153 but not vice versa [13, 16, 17] while the interdependence of Nup153 and TPR for NPC localization is controversial [13, 17]. Similar to MEL28/ELYS, a fraction of the basket nucleoporins Nup50 and Nup153 is found on decondensing chromatin at the early stages of mitotic exit [17, 18]. Nup153 is reportedly not required for mitotic NPC assembly [19, 20]. Whether Nup50 chromatin binding, or Nup50 in general, plays a role in NPC assembly has not been studied so far.

Our current knowledge regarding Nup50 centers around its role as an auxiliary component in nuclear transport (for review see [21, 22]). It can interact with nuclear transport receptors (importin α, importin β, transportin, CRM1) and Ran [10, 15, 16, 23, 24] and has been suggested to act as a platform to support the RanGTP-mediated dissociation of complexes consisting of importin α, importin β and import cargos in a yet ill-defined mechanism. It thus supports the nuclear import reaction, which critically depends on the disassembly of import complexes. A similar accessory role in nuclear transport has been described for the *Saccharomyces cerevisiae* counterpart of Nup50, Nup2 [25–27].

Here we show that, in addition to its role in nuclear transport, Nup50 has a distinct function in mitotic NPC assembly. It binds to and stimulates RCC1’s guanine nucleotide exchange activity towards Ran, which is crucial for mitotic NPC assembly. Interestingly, while this function determines a crucial role for Nup50 in mitotic NPC assembly, this does not require Nup50 to be localized to NPCs.

## Results

### Nup50 is required for nuclear pore complex assembly

To gain insight into Nup50 function we generated antibodies against the Xenopus protein. The antibodies recognize two bands at a size of 50 kDa (Fig. S1A) which can be efficiently depleted from egg extracts (Fig. 1A, S1A). In the egg extract system, nuclear assembly including the formation of a closed NE and NPCs can be faithfully recapitulated [28, 29]. When demembranated sperm-DNA is added as a substrate to the extracts, a nucleus with a functional NE and NPCs is formed (Fig. 1B, untreated and mock). Upon depletion of Nup50 a closed NE, indicated by the smooth membrane staining, forms which, however, lacks NPCs (Fig. 1B, C). This is indicated by the absence of the NPC marker mAB414, which recognizes a subset of FG-nucleoporins not including Nup50 and serves as a robust marker for intact NPCs [30]. Consistent with the absence of NPCs, the nuclei are not competent for nuclear import (Fig. S1B) and, hence, remain of small size. Addition of recombinant Nup50, expressed and purified from bacteria, to approximately endogenous levels (Fig. 1A, S1C) rescues the NPC assembly phenotype (Fig. 1B, C), showing the point specificity of the depletion.

**Figure 1:**
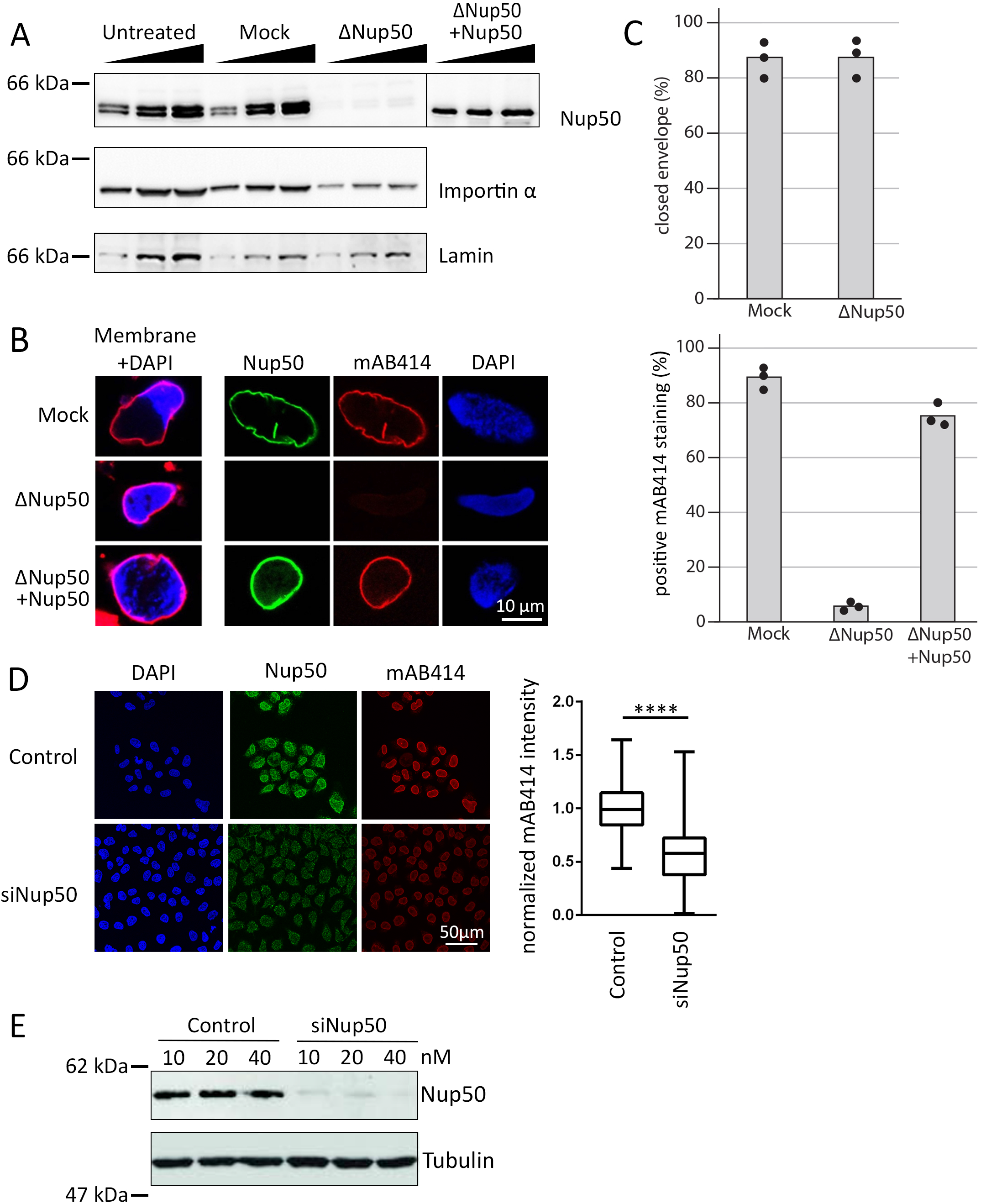
Nup50 is crucial for NPC assembly. (A) Western blot analysis of mock and Nup50 depleted Xenopus egg extracts, with or without addition of recombinant Nup50. 1,2, and 4μl of extracts were analyzed with indicated antibodies. (B) Confocal microscopy images of fixed nuclei assembled for 120 min in mock depleted (mock) and Nup50 depleted (ΔNup50) *Xenopus* egg extracts supplemented with either buffer or recombinant Nup50. In left column, membranes were pre-labelled with DiIC18 (1,1’-Dioctadecyl-3,3,3’,3’-Tetramethylindocarbocyanine Perchlorate, red) and chromatin was stained with DAPI (4′,6-Diamidin-2-phenylindol, blue). Three right columns show the immunofluorescence staining for Nup50 (green), NPCs (mAB414, red) on the chromatin (DAPI). Scale bars: 10 μm. (C) The average percentage of closed nuclear envelopes (upper panel) as well as mAB414 positive nuclei (lower panel) for 100 randomly chosen chromatin substrates in each of 3 independent experiments is shown. Data points from individual experiments are indicated. (D) HeLa cells were transfected with 20 nM control or Nup50 siRNA. 72 h post transfection, the cells were fixed with 4% PFA and stained with antibodies against Nup50 and mAB414, chromatin was labeled with DAPI. Scale bars: 50 μm. The box plots and whiskers show the median, interquartile range and minimum and maximum values of the mAB414 staining intensity of control (n=86) versus Nup50 siRNA treated (n=204) cells. Mann-Whitney test showed a p-value < 0.0001 between the control and siRNA condition. (E) Western blot analysis of HeLa cells treated as in (D) 72 h post transfection.

Thus, in the cell free system Nup50 plays a crucial role in NPC assembly. Next, we tested whether a similar phenotype can be observed in cells. SiRNA-mediated downregulation of Nup50 expression in human HeLa cells (Fig. 1D, E) reduced mAB414 staining, consistent with a role of the protein in NPC assembly. A reduced mAB414 staining has been previously reported upon downregulation of nucleoporins with crucial functions in NPC assembly [31, 32].

It has been previously reported that a Nup50 knock-out mouse is not viable but that fibroblasts derived from mouse embryos can be kept in culture and show an apparent normal mAB414 staining [16]. We speculate that the discrepancy between the human and Xenopus system versus the mouse phenotype could arise because in contrast to humans, rodents like mouse and rats possess two Nup50 related genes. In mouse, aside from the main locus (1700030K07Rik) on chromosome 15, named Nup50A here, a paralogue (1700123L14Rik), named Nup50B here, can be found on the chromosome 6. Although referred to as a pseudogene, this locus is effectively expressed as indicated in the expression atlas [33] and the mouse genome database [34] and has been previously named Nup50rel [16]. We propose that Nup50B might substitute the canonical Nup50 paralogue at least in mouse fibroblasts. Indeed, whereas downregulation of one of the two mouse Nup50 orthologues only moderately reduced mAB414 staining in mouse NIH3T3 cells, the combined application of siRNAs against both targets reduced mAB414 staining (Fig. 2A, B). Please note that the canonical Nup50 paralogue can be detected with antibodies in Western blotting while we could not assay explicitly for Nup50B due to a lack of a specific antibody (Fig. 2C).

**Figure 2:**
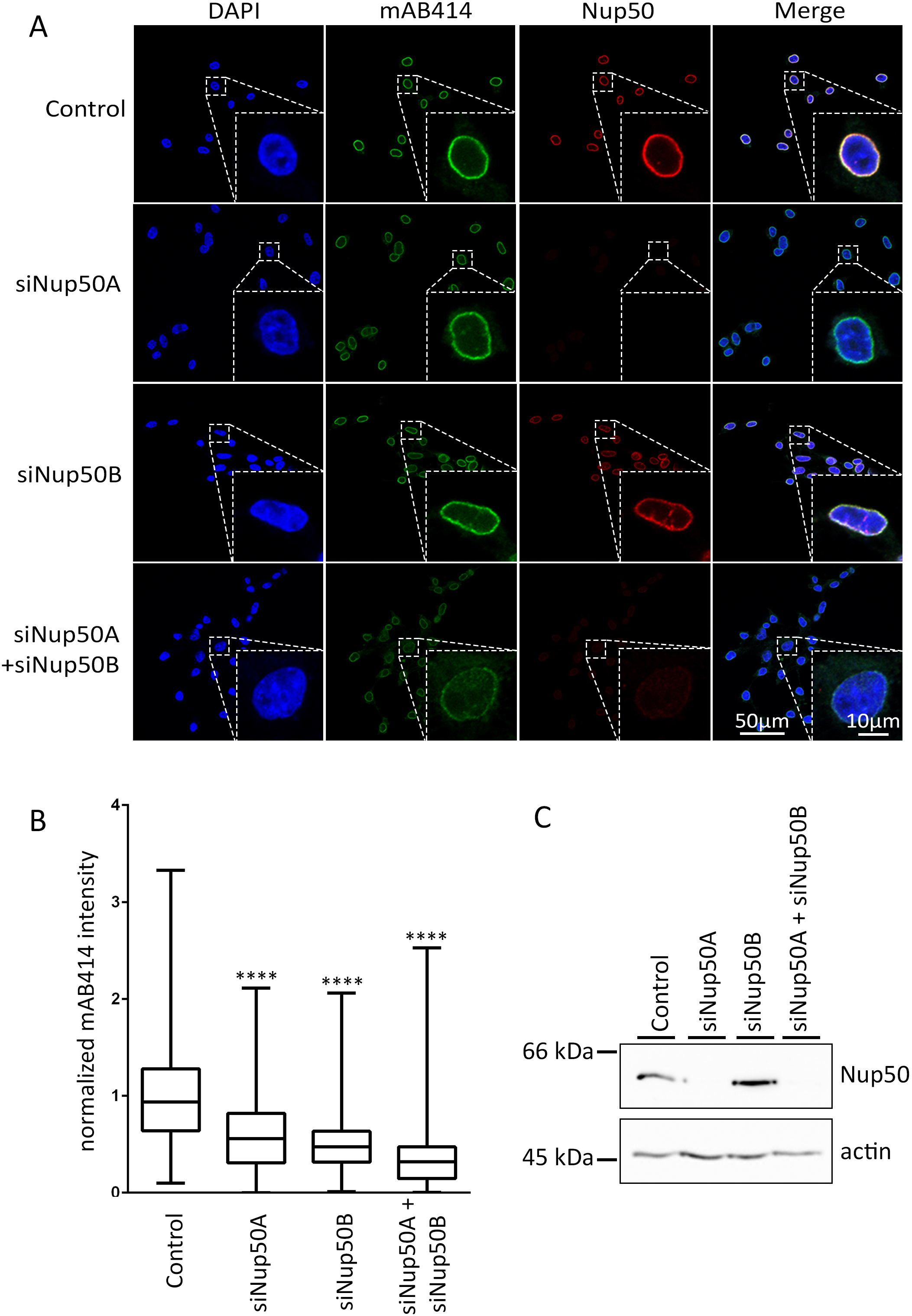
Two Nup50 paralogues in mouse cells. (A) Mouse 3T3-NIH cells were transfected with 20 nM control, Nup50A, Nup50B or a combination of Nup50A and Nup50B siRNA. After 72h cells were fixed and stained with mAB414 (green) and antibodies against mouse Nup50 (red). Chromatin is stained with DAPI (blue). The merge of the three channels is shown on the left column. Scale bar: 50μm. An insert shows a zoom on a representative nucleus for each picture. Scale bar: 10μm. (B) Quantitation of the mAB414 rim intensity relatively to the control. The box plots and whiskers show the median, interquartile range as well as and minimum and maximum value of control (n=551), siNup50A (n=707), siNup50B (n=329) and siNup50A + siNup50B (n=525) conditions. Data were obtained from four independent experiments. Mann-Whitney test showed a p-value < 0.0001 between the control and siRNA conditions. (C) Western blot showing the amount of Nup50A (top row) for each of the experimental condition. The actin (lower row) is shown as a loading control.

Together, these data indicate that Nup50 has a crucial function in NPC assembly, a feature which has been so far overlooked, likely because of the potential redundancy of the two Nup50 paralogues in mice.

### Nup50 interacts with MEL28/ELYS but both act independently in mitotic nuclear pore complex assembly

Interestingly, a similar phenotype, a closed NE without NPCs, has been previously observed upon depletion of the Y-complex [32] and MEL28/ELYS [5, 35], which recruits the Y-complex to the chromatin template during mitotic NPC assembly. Nup50 and MEL28/ELYS interact in Xenopus egg extracts as shown by co-immunoprecipitation (Fig. S2A, B), consistent with a previous report [36]. However, although Nup50 depletion reduces the MEL28/ELYS levels by about 65% and MEL28/ELYS depletion reduces Nup50 by about 15% (Fig. S2C, D), we do not think that the identical phenotypes of Nup50 and MEL28/ELYS is due to a co-depletion of the other factor: The MEL28/ELYS depletion phenotype can be reverted by expression of the protein from the corresponding mRNA [37] and the Nup50 depletion is rescued by addition of recombinant Nup50 (Fig. 1B, C).

Both Nup50 and MEL28/ELYS are recruited early to chromatin during mitotic NPC assembly [18] and in during in vitro nuclear assembly in the Xenopus system (Fig. 3A, B (mock control) and Fig. S3A and B). We therefore tested whether chromatin recruitment of the two proteins depends on each other. Under control conditions, MEL28/ELYS is first found on the chromatin template, about 10 min after initiation of the reaction and later detected at the NE as part of NPCs. In the absence of Nup50, MEL28/ELYS is recruited to chromatin but, consistent with the lack of NPC assembly (see Fig. 1), is not found at the NE at later time points and remains spread on chromatin. Similarly, MEL28/ELYS depletion does not prevent Nup50 chromatin recruitment. However, at later time points Nup50 is not enriched at the NE due to the absence of NPCs, reported for MEL28/ELYS depletion [5, 35] and indicated by the absence of mAB414 staining.

**Figure 3:**
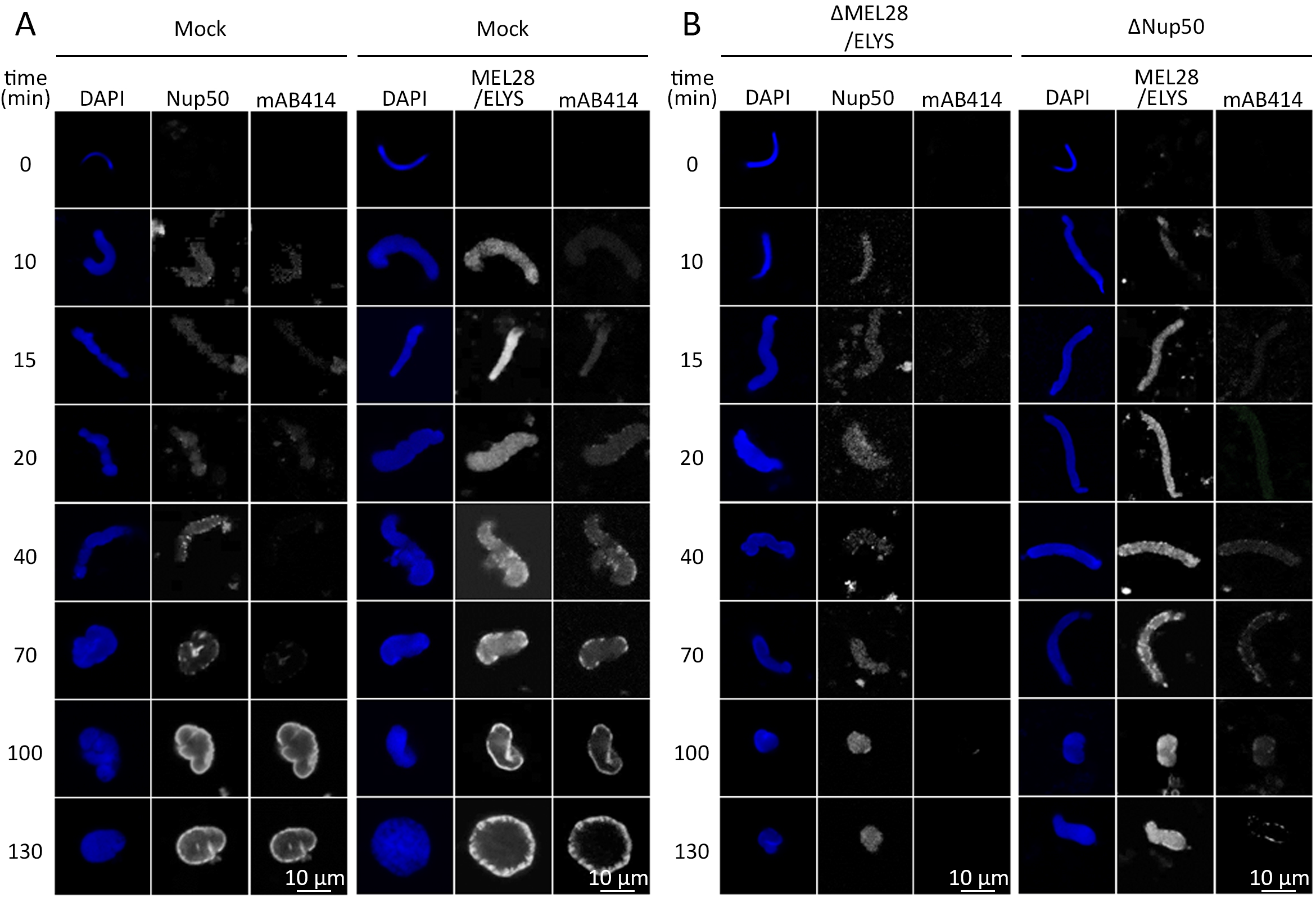
Nup50 and MEL28 do not depend on each other for chromatin binding. (A, B) Demembranated sperm chromatin was preincubated in Xenopus egg extract, depleted for MEL28/ELYS or Nup50 (B) or control treated (Mock, A). After 10 min, membranes were added to the reaction. Reactions were stopped at the indicated time points by fixation and analyzed by confocal microscopy after immunostaining with α-MEL28/ELYS (A), α-Nup50 (B), and mAb414 (A, B). Chromatin was stained with DAPI (blue in the overlay). Scale bar: 10 μm.

### Nup50 binds chromatin via its N-terminal basic motif

During mitotic exit, a relatively substantial subfraction of Nup50, particularly in comparison to other nucleoporins, is found early on the chromatin, approximately 7 min before onset of nuclear import ([18], see also Fig. 3, S3). We therefore tested whether Nup50 can directly interact with chromatin. We generated recombinant EGFP-tagged Nup50 and tested its binding to DNA-coated magnetic beads (Fig. 4). Upon preincubation with egg extracts (right panel), the DNA is rapidly chromatinized [38] whereas in the absence of egg extracts (left panel), pure DNA binding can be detected. In these assays, Nup50 can bind both chromatin and DNA similarly to a C-terminal fragment of MEL28 (aa 2290-2408), which served as a positive control [39].

**Figure 4:**
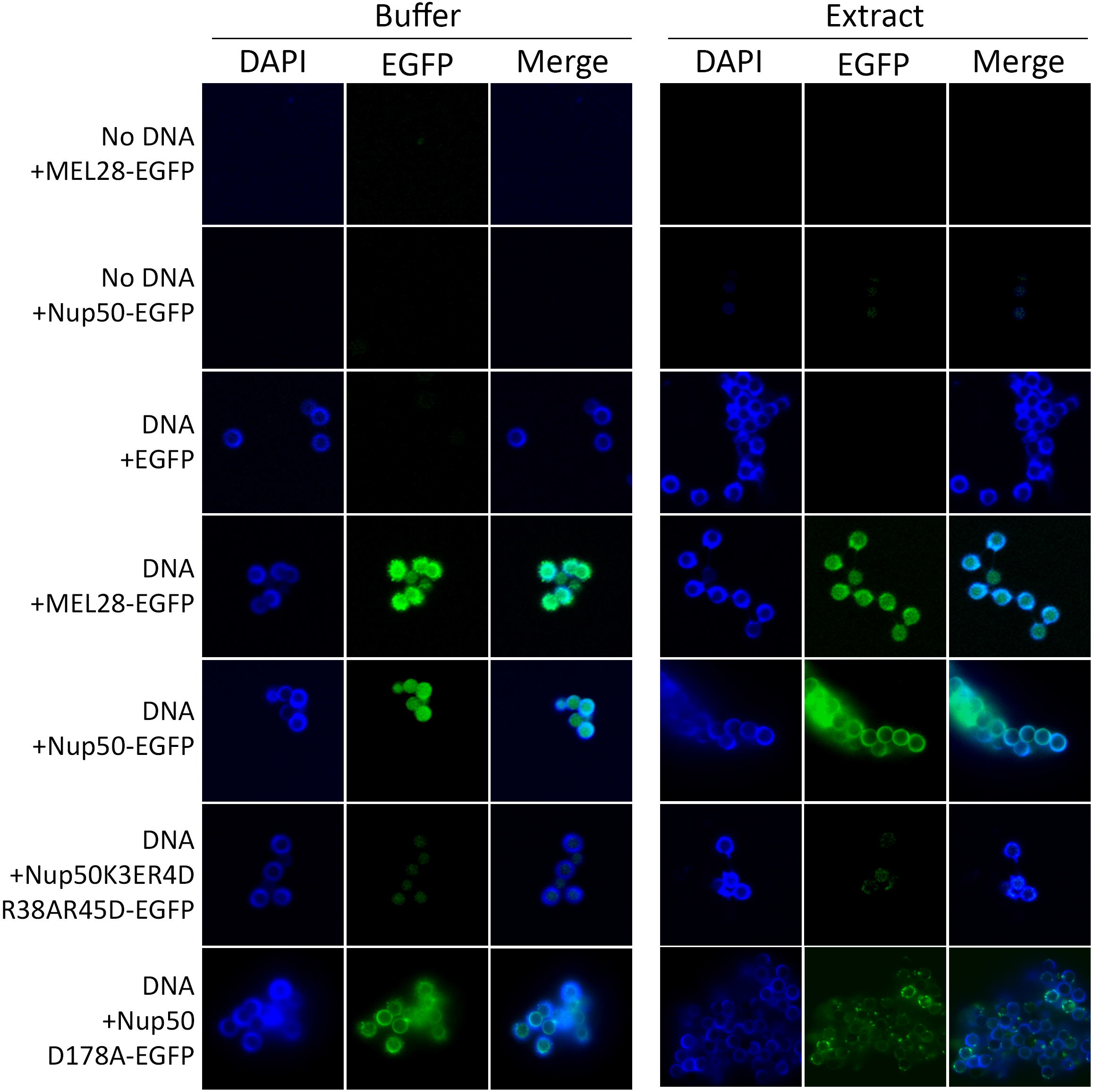
Nup50 is a chromatin binding protein. 3 μM recombinant EGFP, EGFP-tagged MEL28/ELYS (aa 2290–2408) or EGFP-tagged Nup50 were incubated with empty or DNA-coated magnetic beads, which were chromatinized with Xenopus egg extracts (right panel) or unchromatinized DNA-beads (left panel). After 3h the beads were re-isolated, washed, co-stained with DAPI and analyzed by confocal microscopy.

Nup50 contains an N-terminal bipartite nuclear localization signal [23, 24]. These motifs often serve, in addition to importin α binding, as chromatin interaction domains [40]. Indeed, mutations in this positively charged region (K3E,R4D,R38A,R45D), known to prevent importin α interaction [24], abolishes chromatin/DNA binding of Nup50 (Fig. 4).

### A short 46 residues-long region of Nup50 is required for its nuclear pore complex localization

To gain insight into Nup50 function in NPC assembly we aimed to identify the parts of the protein critical for its NPC localization. Nup50 is comprised of an N-terminal importin α/chromatin interaction domain and a C-terminal Ran binding domain [23]. Using EGFP-tagged fragments of Nup50, expressed in HeLa cells, we found that neither domain is strictly required for the localization of Nup50 to NPCs (Fig. 5A). Further truncations yielded a minimal 46 amino acid region (aa 144-189) required for NPC localization. This region of Nup50 shows a higher evolutionary conservation than other parts of the proteins (Fig. 5B); that is 80.4% sequence identity between the Xenopus and human proteins versus 59.6% for the entire protein. Several residues in this domain are highly conserved and for a number of them, if mutated in the context of the full-length protein, NPC localization was abolished (Fig. 5C). By pulldown assays, we identified the minimal NPC binding region of Nup50 as a domain binding to both MEL28/ELYS and Nup153 (Fig. 5D). In the context of this minimal fragments the mutations that abolish NPC binding prevented the interaction to both MEL28/ELYS and Nup153 (Fig. 5D). This minimal region is included in the MAR domain of *S.cerevisiae* Nup2 [41] which was identified as the minimal fragment rescuing Nup2 meiotic function and its NPC localization. An alignment of *S.cerevisiae* MAR domain with metazoan Nup50 shows a better alignment with the 144-189 region than the rest of the protein (Fig. S4). Consistently, sequence identity between *X. laevis* 144-189 and *S.cerevisiae* MAR domain is 32.6% versus 22.7% for the full length proteins.

**Figure 5:**
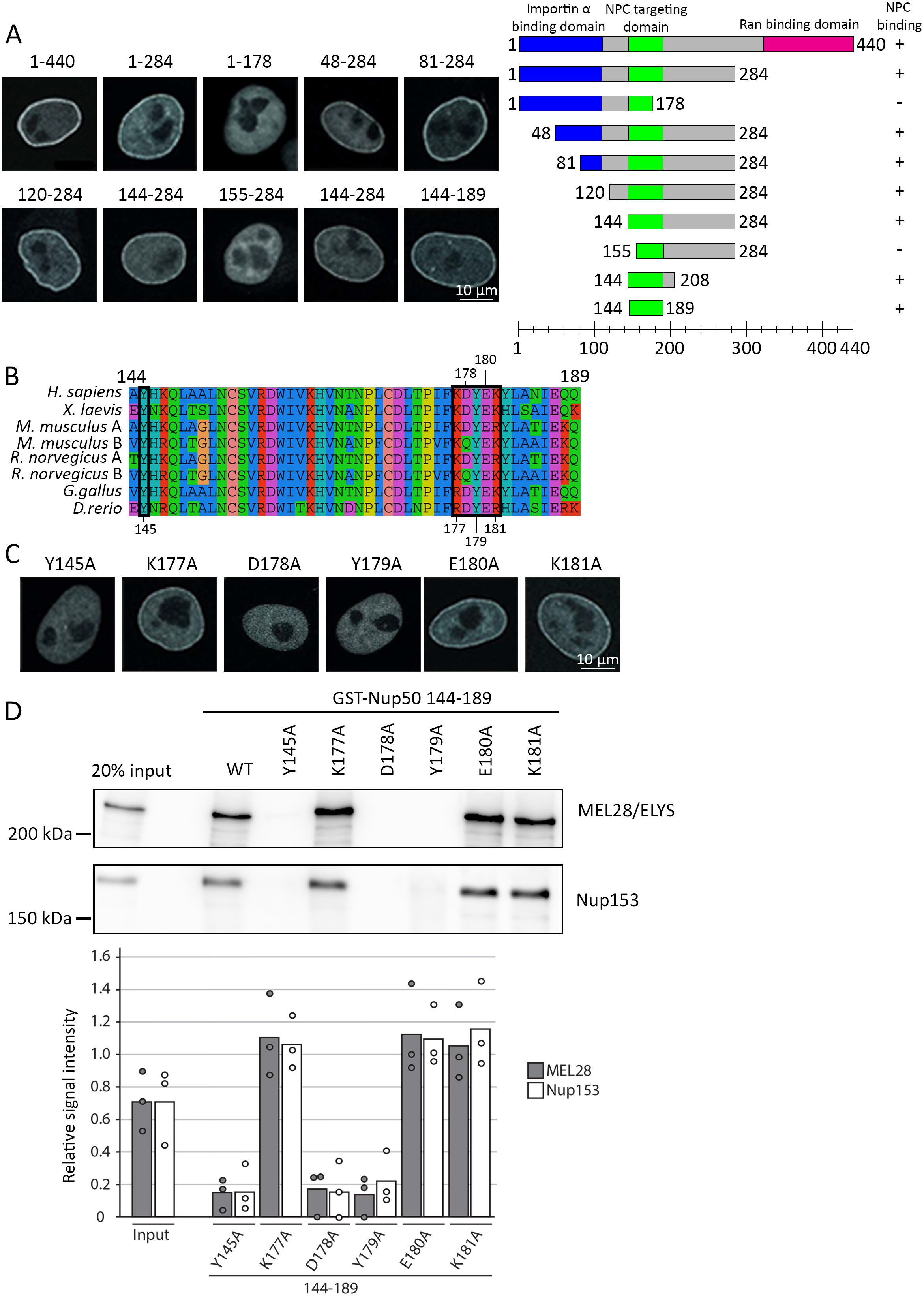
A minimal 46aa region is required for Nup50 NPC localization. (A) HeLa cells were transfected with EGFP-Nup50 and different truncations. After 24h cells were shortly pre-treated with 0.1% Tx100 in PBS, fixed with 4% PFA and analyzed by confocal microscopy (left panel). Scale Bar: 10 μm. The graph (right panel) shows the Nup50 domains (blue: Importin α interaction domain, green: NPC targeting domain and pink: Ran binding domain) and the truncation tested. Ability to bind the nuclear rim is indicated on the right. (B) Sequence alignment of Nup50 144–189, species are indicated on the left, the black boxes highlight residues tested by single point mutation in (C) and (D), the residues number are shown above or below the alignment. The color scheme indicates the type of amino acids according to the alignment software default setting. (C) HeLa cells were transfected with EGFP-Nup50 comprising different point mutations in the minimal NPC binding region and analyzed as in (A). Scale Bar: 10 μm. (D) GST-fusion constructs of the Xenopus Nup50 minimal NPC binding fragment (aa 144–189) comprising no or single point mutations were incubated with Xenopus egg extracts. Starting material as well as bound proteins were analyzed by Western blotting with indicated antibodies. The quantitation shows the average MEL28 and Nup153 bead bound signal from three independent experiments, normalized to the wild type signal. Data points from individual experiments are indicated.

### A minimal N-terminal fragment is required for nuclear pore complex assembly independent of NPC localization

Having identified crucial amino acids for Nup50 binding to MEL28/ELYS and Nup153, which are also required for NPC localization, we wondered whether these interactions are crucial for Nup50 function in NPC assembly. We used the Xenopus egg extract system to deplete Nup50 and add back the wild-type or different mutant versions. Surprisingly, the mutants which abolished NPC localization rescued the NPC assembly phenotype (Fig. 6A, B). Despite the rescue and consistent with our data for human cells, these mutants did not localize to NPCs (Fig. S5). Also, a Nup50 version which does not interact with importin α/chromatin (K3E,R4D,R38A,R45D) or directly with importin β (FGYG, where all five FG sequences in the Xenopus protein have were replaced by YG) was able to replace the wild-type protein in NPC assembly indicating that these specific protein-protein interactions are not crucial for this function.

**Figure 6:**
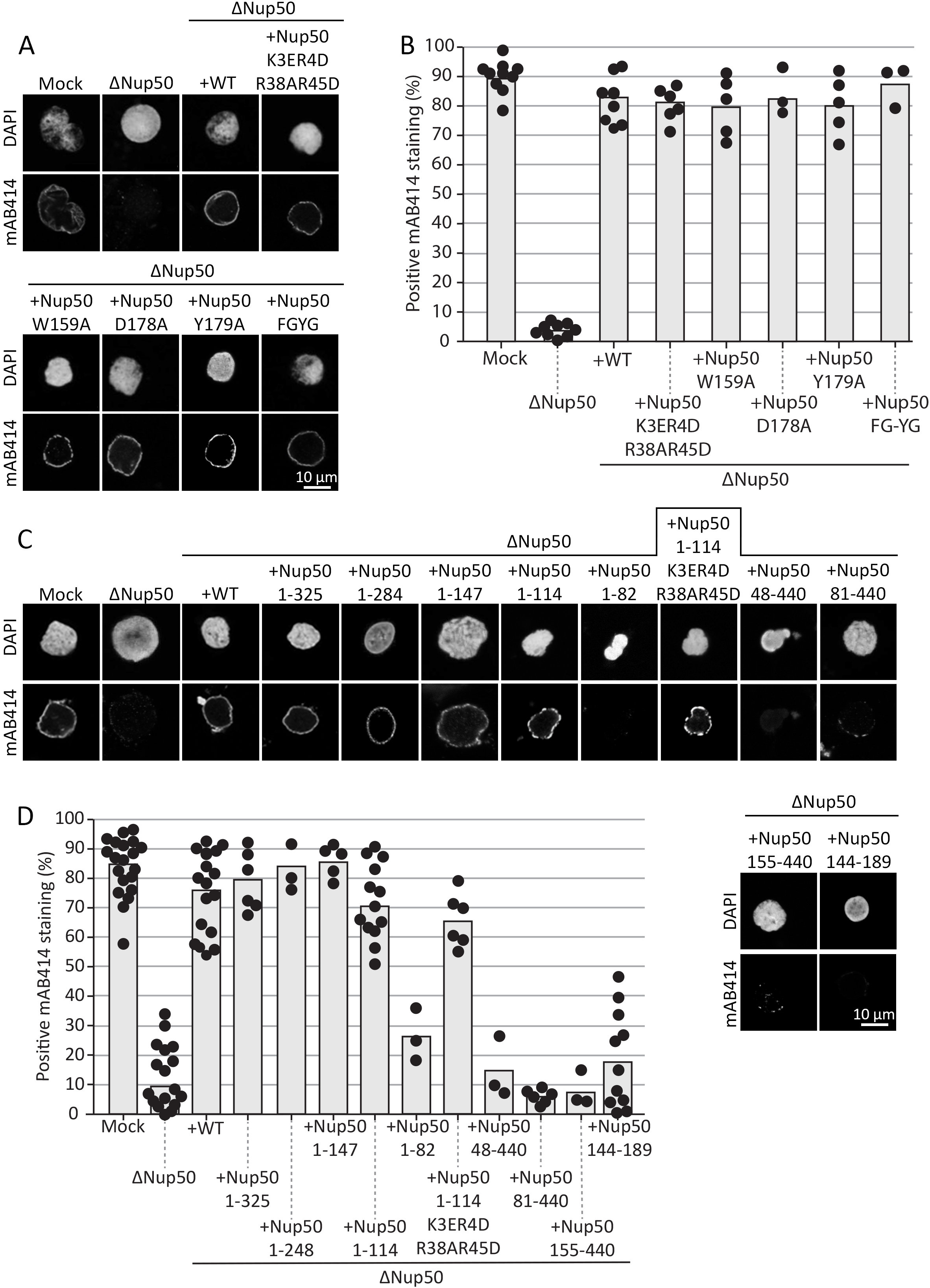
A N-terminal Nup50 fragment is required for NPC assembly. (A) Confocal microscopy images of nuclei assembled for 120 min in mock depleted, Nup50 depleted (ΔNup50), and Nup50 depleted *Xenopus* egg extracts supplemented with recombinant wild type Nup50 or different mutants. Nuclei were fixed in 4% PFA and 0.5 % glutaraldehyde, stained for NPCs (mAB414) and the chromatin (DAPI). Scale bar: 10 μm. (B) Average percentage of mAB414 positive nuclei for 100 randomly chosen chromatin substrates in each of at least three independent experiments shown in (A). Data points from the individual experiments are indicated. (C) Confocal microscopy images of nuclei assembled with N-terminal Nup50 truncations and an minimal N-terminal fragment. Samples were analyzed as in (A). (D) Average percentage of mAB414 positive nuclei for 100 randomly chosen chromatin substrates in each of at least three independent experiments shown in (C). Data points from the individual experiments are indicated.

To identify the regions of Nup50 crucial for NPC assembly, we employed different Nup50 fragments in the cell-free assay. These experiments revealed that a minimal fragment comprising aa 1-114 is sufficient to substitute full-length Nup50 in NPC assembly (Fig. 6 C, D). Thus, neither the Ran binding domain, located in the C-terminus of the protein, nor MEL28/ELYS, Nup153, importin α and importin β binding of Nup50 are required for the protein’s function in NPC assembly, consistent with the above data.

### Nup50 binding to membranes is not required for nuclear pore complex assembly

Nup2, the *S. cerevisiae* orthologue of Nup50, can reportedly bind liposomes via a pleckstrin homology (PH) domain [42]. We tested whether vertebrate Nup50 could similarly bind liposomes. Indeed, EGFP-tagged Nup50 can bind to large liposomes, giant unilamellar vesicles (Fig. S6). Floatation experiments with small liposomes (size range 30-400 nm) confirmed these data. We were unable to map the membrane binding domain within Nup50 as truncation from the N-as well as from the C-terminus abolished membrane interaction. However, as the same truncations were functional in NPC assembly, this observation indicates that the Nup50 membrane interaction is dispensable for its function in NPC formation.

### Both Nup50 orthologues are functional in nuclear pore complex assembly

We speculated above that the mouse Nup50 paralogues could functionally substitute each other in NPC assembly. To further test this hypothesis, we replaced Nup50 in egg extracts by either of the two mouse orthologues, recombinantly expressed in bacteria (Fig. 7A). Interestingly, both orthologues could rescue the NPC assembly phenotype (Fig. 7B,C). However, whereas Nup50A localized both to chromatin and the NE in the assembly reaction, Nup50B was only found on chromatin.

**Figure 7:**
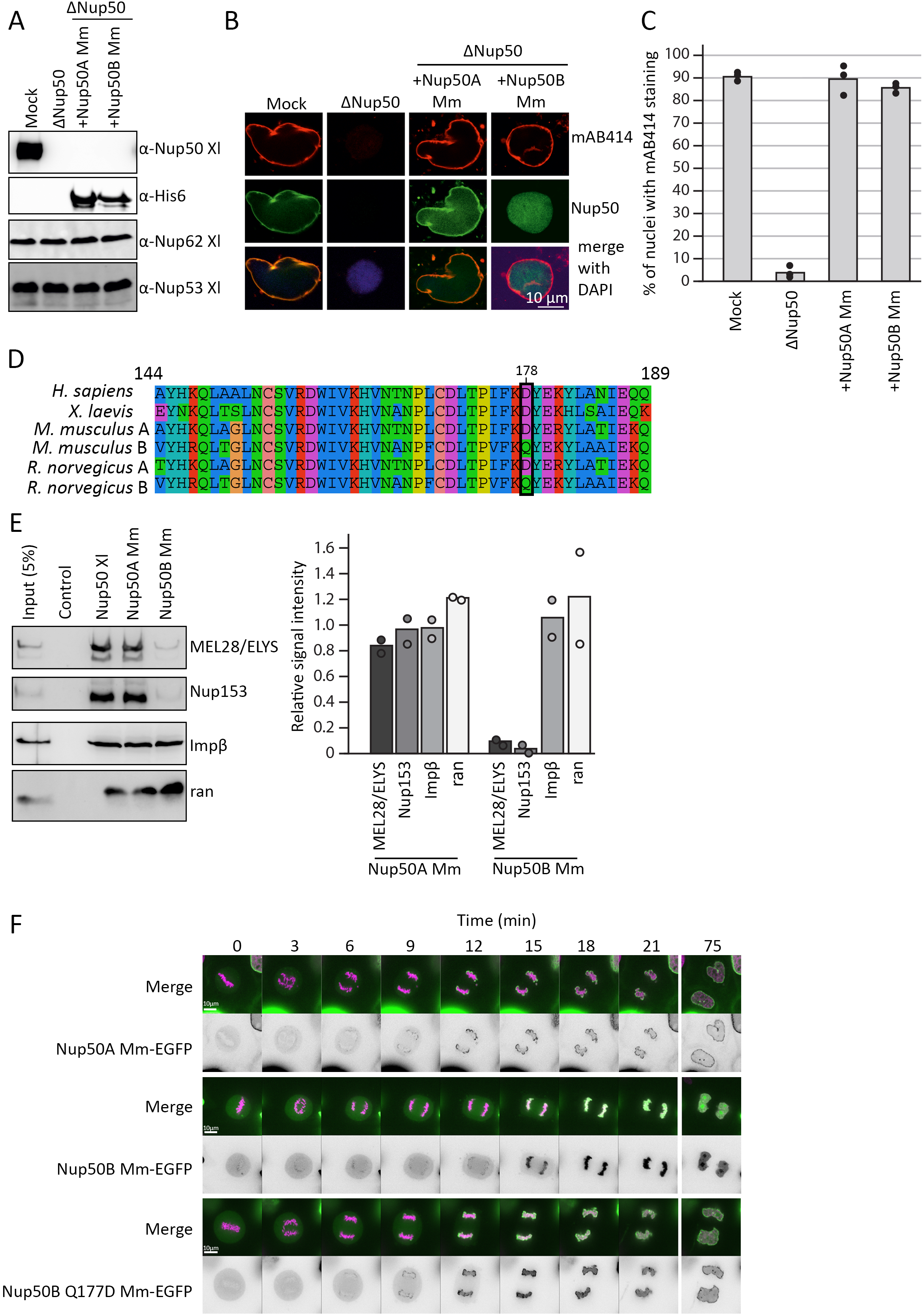
Both mouse Nup50 orthologues can replace the Xenopus protein. (A) Western blot analysis of mock and Nup50 depleted Xenopus egg extracts (ΔNup50), with or without addition of recombinant mouse Nup50 orthologues. Samples were analyzed with Xenopus Nup50 antibodies, which does not recognize the mouse proteins, which in turn were detected with an His6-antibody. (B) Confocal microscopy images of fixed nuclei assembled for 120 min in mock depleted (mock) and Nup50 depleted (ΔNup50) *Xenopus* egg extracts supplemented with recombinant mouse Nup50 orthologues. Nuclei were stained for Nup50 (Nup50 antibody for Xenopus protein, His6 for mouse protein) and NPCs (mAB414, red) on the chromatin (DAPI). Scale bar: 10 μm. (C) The average percentage of mAB414 positive nuclei for 100 randomly chosen chromatin substrates in each of three independent experiments (performed as in B) is shown. Data points from individual experiments are indicated. (D) Sequence comparison of the conserved 46aa fragment of the human, mouse, rat and *Xenopus laevis* sequences, with the residue 178 (Xenopus numbering), crucial for NPC binding in the Xenopus protein is highlighted (black box). The color scheme indicates the type of amino acids according to the alignment software default setting. (E) GST-fusion constructs of the Xenopus Nup53 RRM domain (aa 162–267, control) full-length Xenopus Nup50 and the two mouse Nup50 orthologues were incubated with Xenopus egg extracts. Starting material as well as bound proteins were analyzed by Western blotting with indicated antibodies. The right panel shows the average MEL28/ELYS, Nup153, importin β and ran bead bound signal from two independent experiments normalized on the Xenopus signal. Data points from individual experiments are indicated. (F) HeLa cells were transfected with EGFP-tagged constructs of both mouse Nup50 orthologues and the Nup50B Q177D (orthologue to the position 178 in Xenopus) mutant. After 24h, the cells were analyzed by life cell imaging. Panel shows cells exiting mitosis (time normalized to metaphase-anaphase transition). The merge shows histone 2B in pink and Nup50 in green, the black and white panel shows Nup50-EGFP signal.

A sequence comparison indicates that within the conserved region identified in the Xenopus protein as minimal NPC binding domain, several amino acid changes occur between the two mouse orthologues (Fig. 7D), including an aspartic acid (D) to glutamine (Q) exchange in the position 177 (corresponding to the position 178 in Xenopus) which is critical for NPC binding in the Xenopus protein. Indeed, GST pulldowns showed that the canonical Nup50A can interact with Nup153 and MEL28 whereas mouse Nup50B shows only a very reduced interaction levels (Fig. 7E).

Transfection of HeLa cells with EGFP-tagged versions of both mouse Nup50 orthologues similarly revealed that Nup50A localized to chromatin and the NE, consistent with published data [43], whereas Nup50B was only found on the chromatin (Fig. 7F). A single point mutation, Q177D in Nup50B, was sufficient to relocalize the EGFP fusion to the NE. Together, these data demonstrate that both Nup50 orthologues can function in NPC assembly and support the notion that NPC localization of Nup50 is not required for its role in mitotic NPC re-assembly.

### A minimal N-terminal fragment interacts with RCC1

To define the function of the N-terminal part of Nup50, crucial for its function in NPC assembly, we performed pulldown experiments of the corresponding human FLAG-tagged fragment from HEK cells. By mass spectrometry several interaction partners were identified and confirmed by western blotting (Fig. 8A, B), including importin α, RCC1 and ANP32A. Whereas ANP32A interacted with human and Xenopus Nup50 in pulldown assays, it only bound to the second mouse Nup50 paralogue, Nup50B (Fig. 8C). As both paralogues can rescue Nup50 depletion in egg extracts it is unlikely that the ANP32A interaction is crucial for the role of Nup50 in NPC assembly. Similarly, the interaction with importin α is unlikely to be crucial for Nup50’s function in NPC assembly as a recombinant Nup50 protein defective in importin α binding can effectively replace endogenous Nup50 in the cell free assays (Fig. 6 C, D).

**Figure 8:**
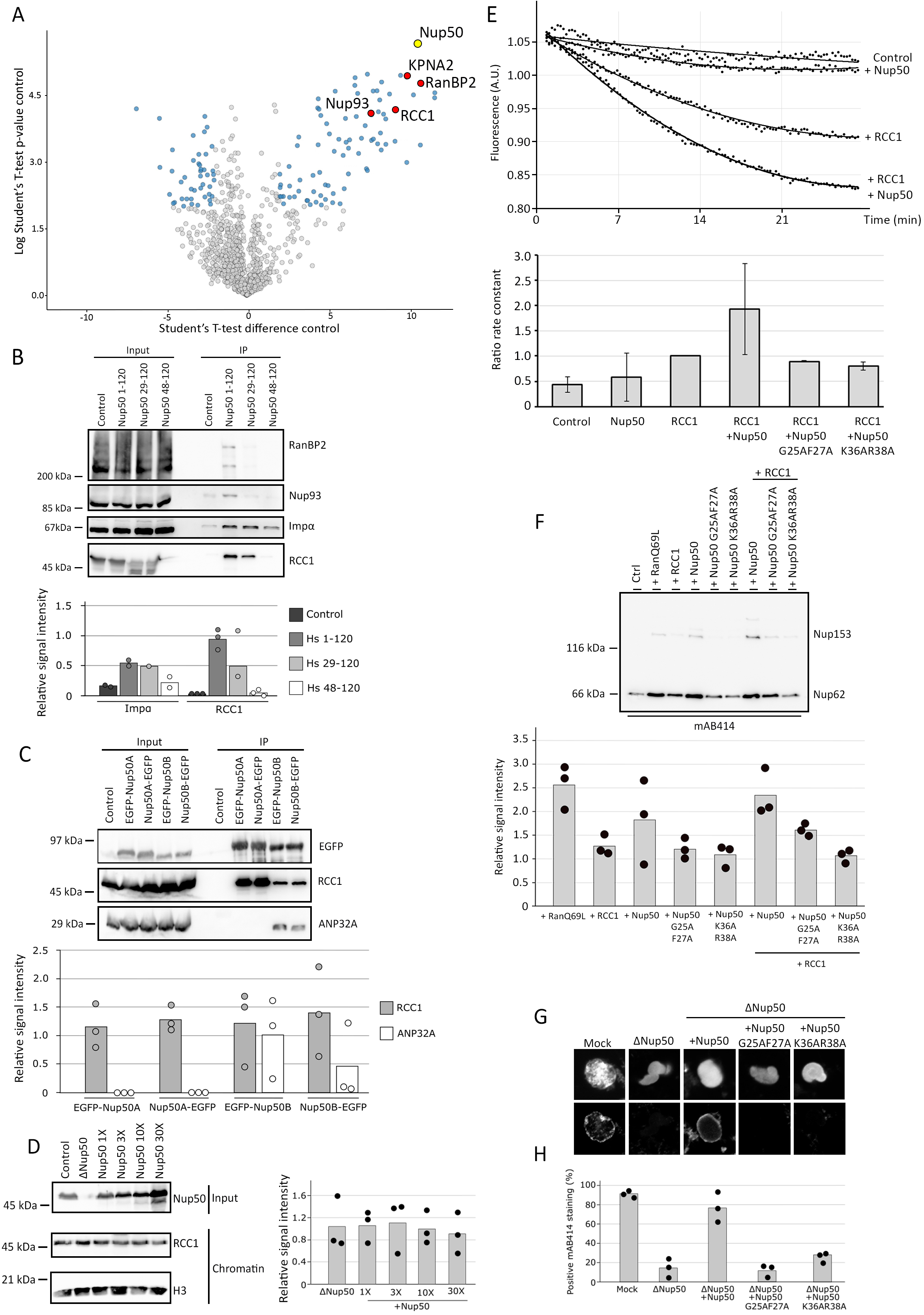
Nup50 stimulates RCC1 activity for NPC assembly. (A;B) HEK293T cells were transfected with empty FLAG-tag vector as a control or FLAG-Nup50 N-terminal fragment (aa 1–120 of the human sequence). 24 h post transfection cells were lysed, FLAG-tagged proteins immunoisolated and analyzed by mass spectrometry (A) with the Volcano blot showing the identified interactors (red) of Nup50 (yellow, data from three independent experiments) and by western blotting (B), with 10% of the inputs loaded. The quantification of the western blot (B, lower panel) shows the mean signal intensity normalized on the input of at least two independent experiments. Data points from individual experiments are indicated. KPNA2 is the gene name of importin α. (C) HEK293T cells were transfected with both Nup50 mouse orthologues N- and C-terminally tagged with EGFP. 24h post transfection cells were lysed, EGFP-tagged proteins immunoisolated and analyzed by western blotting. The quantification of the western blot (C, lower panel) shows the mean signal intensity normalized on EGFP of three independent experiments. Data points from individual experiments are indicated. (D) Xenopus sperm chromatin (6 000 sperm heads/μl were incubated with 120 μl of control, Nup50 depleted or egg extracts supplemented with excess (1X = 0,07μM final) recombinant Nup50 and re-purified. Total input extracts were analyzed by western blotting against Nup50 and isolated chromatin were analyzed by western blotting against RCC1 and Histone H3 as a loading control (left panel). The quantification (right panel) shows the mean signal intensity of the RCC1 signal normalized over Histone H3 from three independent experiments. Data points from individual experiments are indicated. (E) 2 μM recombinant Ran, loaded with MANT-GDP was incubated with 2 mM GppNHp in buffer control, supplemented with 2 nM recombinant RCC1, 20 nM recombinant Nup50 proteins, or RCC1 and Nup50 together. GDP to GppNHp exchanged was monitored by the decrease in MANT fluorescence of the liberated GDP-MANT. Quantitation shows the ratio of the rate constant normalized in the Ran + RCC1 + GppNHp condition from five independent experiments. (F) Xenopus egg extract were supplemented with 5 μM RanQ69L, RCC1, Nup50 wild type and RCC1 binding mutants as well as a combination of 5 μM Nup50 and 5 μM RCC1. After 90 min, annulate lamellae were isolated by centrifugation and quantified by western blotting with mAB414 antibody. Quantitation shows the relative Nup62 signal as a mean from four independent experiments, normalized to the buffer control. Individual data points are indicated. (G) Confocal microscopy images of nuclei assembled for 120 min in mock depleted, Nup50 depleted (ΔNup50), and Nup50 depleted *Xenopus* egg extracts supplemented with recombinant wild type Nup50 or RCC1 binding mutants. Nuclei were fixed in 4% PFA and 0.5 % glutaraldehyde, stained for NPCs (mAB414) and the chromatin (DAPI). Scale bar: 10 μm. (H) Average percentage of mAB414 positive nuclei for 100 randomly chosen chromatin substrates in each of at least three independent experiments shown in (G). Data points from the three individual experiments are indicated.

The guanine nucleotide exchange factor (GEF) of Ran, RCC1, reportedly plays a crucial role in NPC assembly [8, 44]. RCC1 dynamically interacts with chromatin and several factors are known to regulate this interaction. As Nup50 also interacts with chromatin, we speculated that it might affect RCC1 chromatin binding. However, RCC1 loading on re-isolated sperm chromatin after incubation in egg extracts was neither affected by Nup50 depletion nor by the addition of excess of Nup50 (Fig. 8D). Alternatively, Nup50 might affect RCC1’s GEF activity. Indeed, in GDP to GTP exchange assays, recombinant Nup50 enhanced RCC1 activity towards Ran by twofold (Fig. 8E). Thus, Nup50 might function in mitotic NPC assembly by stimulating RCC1 activity, which in turn generates RanGTP in the vicinity of chromatin as crucial factor for NPC formation. Ectopic addition of a high concentration of RanGTP or of the constitutively active mutant RanQ69L to egg extracts in the absence of chromatin, stimulates formation of annulate lamellae (AL) [44], membrane stacks with tightly packed NPCs in the ER-like membranes. Interestingly, addition of Nup50 also increases AL formation under these conditions and co-addition of RCC1 increases the effect (Fig. 8F). As AL form in the absence of chromatin this finding is consistent with the notion that Nup50 does not act on RCC1 by recruiting it to chromatin but rather stimulates its enzymatic activity.

To directly assess the contribution of the RCC1-Nup50 interaction to NPC assembly, we generated Nup50 mutants defective in RCC1 binding without affecting the importin α interaction (Fig. S7A). The corresponding Xenopus proteins, Nup50 G25A,F27A and Nup50 K36A,R38A, failed to stimulate RCC1 GEF activity (Fig. 8E). Accordingly, neither protein could rescue depletion of endogenous Nup50 in the *in vitro* nuclear assembly assay (Fig. 8 G, H). Similarly, addition of either mutant alone or in combination with RCC1 could not stimulate AL formation in egg extracts (Fig. S7B).

## Discussion

Here we show that the nucleoporin Nup50 plays a crucial role in mitotic NPC assembly. Surprisingly, NPC localization of Nup50 is not required for its function in NPC assembly. Consistent with this conclusion, Nup50 is reportedly absent in NPCs of annulate lamellae [17], ER membrane stacks in which NPCs have integrated at a distance to chromatin [45]. In addition, Nup50 has a short residence time at the NPC [46] and is continuously exchanged at NPCs even in non-dividing cells [47], two features that are typical for peripheral nucleoporins that are not part of the NPC scaffold. Our data suggest that the N-terminus of Nup50 binds to the RanGEF RCC1 and stimulates its GDP-GTP exchange activity towards Ran.

In addition to its crucial role in nuclear transport, Ran has important functions during mitosis. It promotes spindle assembly during mitotic entry and NPC re-assembly during mitotic exit (for review see [7]). The chromatin-bound RanGEF RCC1 is crucial for the latter function. It generates a high RanGTP concentration around chromatin which is required to release essential spindle assembly factors such as Tpx2 or NuMA from inhibitory nuclear transport receptors. Although the precise RanGTP targets functioning in NPC assembly are currently unknown, a similar function has been proposed for NPC formation. Our data indicate that the interaction of Nup50 with RCC1 is crucial for its function in mitotic NPC assembly. A short N-terminal fragment interacting with RCC1 and stimulating its RanGEF activity is sufficient to replace the endogenous Nup50 in *in vitro* NPC assembly assays (Fig. 6). Accordingly, Nup50 mutants that cannot bind and activate RCC1, are unable to substitute the wild type protein (Fig. 8). Stimulation of RCC1 should increase RanGTP production in the proximity of chromatin and promote NPC assembly. Several additional factors are known to modulate RCC1 activity, including chromatin binding, which stimulates RCC1 [48], and RanBP1, which inhibits RCC1 by binding and preventing its chromatin interaction [49]. With Nup50, we now identify another regulator of RCC1 and the Ran system important for NPC re-assembly during mitotic exit.

Notably, we attempted but failed to compensate for Nup50 depletion in the Xenopus extract system by addition of RCC1 or RanQ69L as appropriate titration of this system presents a significant challenge. Addition of RanQ69L is known to lead to ectopic NPC assembly in AL and block NPC assembly [44]. Furthermore, RCC1 addition might not be sufficient to substitute for the loss of Nup50 since RanBP1, which is highly abundant in egg extracts, blocks RCC1 activity [49, 50].

Much like MEL28/ELYS, a fraction of the nucleoporin Nup50 pool is found on decondensing chromatin at the early stages of mitotic exit [17, 18]. This early localization is consistent with our finding that Nup50 can bind chromatin in *in vitro* assays (Fig. 3 and 4). Nevertheless, chromatin binding of Nup50, at least in the in vitro nuclear assembly reactions, is not crucial for its function in NPC assembly. Later in the nuclear reassembly process, a larger fraction (about 80%) of Nup50 associates with NPCs in human cells [18] and in the *in vitro* Xenopus assemblies (Fig. 3). At later time points a more intensive rim staining is observed for Nup50. This NPC localization depends on NPC formation because in the absence of MEL28/ELYS, which reportedly blocks NPC assembly, the rim staining is apparently lost (Fig. 3B). We speculate that the first pool of Nup50 recruitment is required for RCC1 stimulation whereas the second pool integrates into assembled NPCs.

Nup50 is localized at the nucleoplasmic side of the NPC [10]. Consistent with this localization, it interacts with Nup153 [15, 16, 51], which is similarly found in this region of the NPC. Down-regulation of Nup153 abolishes the rim but not the nucleoplasmic localization of Nup50 indicating that it is no longer found at NPCs [13, 17, 51, 52]. Interestingly, loss of Nup153, which leads to Nup50 displacement from the NPC, does not affect assembly of the core structure of the NPC [17], which is consistent with our finding that Nup50’s NPC localization is not crucial for its role in NPC assembly. We identified a conserved 46 amino acid region of Nup50 that is required for NPC localization an is a Nup153 and MEL28/ELYS binding domain. This minimal region is included in the Nup50 fragment (aa 1-214 of the human sequence) previously identified as Nup153 interaction site [15]. It is also required for MEL28/ELYS interaction, which has been identified as Nup50 interaction partner in egg extracts [36]. We have not been able to separate the Nup153 and MEL28/ELYS binding sites and as all NPC binding mutants affect both interactions similarly, we suspect that both proteins use indeed the same sequence motifs for interaction with Nup50. In the light of the published data from human cells, it is likely that Nup153 binding is the determinant for Nup50’s NPC localization. Nup50 can also interact in pulldown experiments with Nup153 via importin α, an interaction requiring its N-terminal importin α binding domain [15] (our unpublished data). However, at least in cell transfection assays, this interaction is not able to recruit Nup50 to NPCs (Fig. 5).

Nup50 is able to bind liposomes (Fig. S6). Interestingly, the *S. cerevisiae* orthologue Nup2 has been suggested to contain a pleckstrin homology domain, a common phosphoinositide binding motif, which correspondingly binds with PI(4,5)P2 containing liposomes [42]. Although we did not observe a phosphoinositide dependence for full-length Xenopus Nup50 membrane binding, this adds Nup50 to the list of nucleoporins that can directly interact with the nuclear membranes. However, at least for NPC interaction and in *in vitro* NPC assembly, membrane binding seems not to be required.

Homozygous Nup50 knock-out mice die during late embryonic development [16], indicating a crucial function of Nup50. The embryos are growth retarded and show neural tube defects. Mouse embryonic fibroblasts isolated from Nup50 null embryos exhibit no obvious defects in cell cycle control or NPC function and nuclei possess normal mAB414 staining. These findings are surprising in light of our experiments. They may suggest that a Nup50-related gene takes over Nup50 function [16]. Genome and EST database searches suggest that a rodent-specific gene, which shows 58,2% and 59,3% amino acids identity in mouse and rat, respectively, and possesses the same domain arrangement as mouse/rat canonical Nup50 (Fig. 2), could compensate for the loss of Nup50. As previously noted [16], this gene is highly expressed in testis but additionally shows detectable expression levels in other tissues [33, 34]. In line with this interpretation, our RNAi experiments in HeLa cells show a marked loss of mAB414 staining upon Nup50 depletion, which is only recapitulated in mouse NIH3T3 cells if siRNAs against both potential orthologues are employed. All tested Nup50 antibodies are directed only against the canonical Nup50A orthologue and, unfortunately, did not detect the second orthologue. Using the auxin inducible degron system, it was recently shown that Nup50-depleted DLD-1 cells are viable and continued to grow, albeit somewhat more slowly than the parental cells [13]. However, in these cells, RCC1 is expressed as fusion with a fluorescent protein and the ubiquitin ligase. This might result in expression or functional changes of RCC1 which could compensate for the loss of Nup50.

The two Nup50 paralogs described in rodents are encoded on different chromosomes and distinct form the two reported human Nup50 isoforms, Npap60L (aa 1-469) and Npap60S (aa 29-469) [53, 54] that are generated by alternative splicing. Npap60L is widely expressed from vertebrates to yeast and corresponds here to Nup2. Npap60S has been only found in humans. Npap60L and Npap60S have been suggested to differentially regulate nuclear import [55]. While Npap60L promotes cargo release from importin α, Npap60S would rather stabilize the importin α-cargo complex. It should be noted that a 29-120 fragment of human Nup50 binds RCC1, arguing that both isoforms could in theory bind RCC1 (Fig. 8B). Additionally, in *Xenopus laevis* two paralogs of Nup50 are found but this is also different from the situation observed in mouse. The two Xenopus paralogs arise from an ancestral whole genome duplication [56] and both loci have retained their exon/intron organization [57]. Moreover, the two paralogues show a 89.5% of sequence identity in Xenopus as compared to approximately 60% in rodents. In Xenopus, both paralogs can be detected by Western blotting depending on the total protein amount loaded and electrophoresis conditions (see Fig. 1A, S2) and are both efficiently depleted by our Nup50 antibodies. So far, we have no data indicating that the Xenopus paralogs could have different biological functions.

Although the Nup50 orthologue Nup2 is not essential in budding and fission yeast, it is vital in the fungus *Aspergilllus nidulans* [58, 59]. During semi-open mitosis, where NPCs only partially disassemble, Nup2 re-localizes to the mitotic chromatin. In anaphase and telophase, Nup2 attracts other nucleoporins to chromatin thereby serving as bridge between the segregating chromatin and NPCs [60]. This prevents loss of NPCs from the NE during nuclear division and guarantees nuclear transport capabilities in G1 phase. Thus, both vertebrate Nup50 and *Aspergillus nidulans* Nup2 secure appropriate NPC numbers and transport competence of the nucleus during mitotic exit, yet by different mechanism. While for yeast Nup2 both chromatin association and interaction with NPCs matter, vertebrate Nup50 does not crucially rely on these capabilities for its function in NPC reassembly.

In summary, here we characterize Nup50, which has been so far mainly described as a nucleoporin enhancing nuclear transport efficiency by interactions with the importin α family and Ran, as factor involved in mitotic NPC re-assembly by enhancing RCC1 activity. This same mechanism of RCC1 activation might also contribute to the function of Nup50 in nuclear transport. Generation of RanGTP by RCC1 is critical for highly efficient and directed transport across the NPC barrier. RCC1 and the Ran system also plays a crucial role early in mitosis, e.g. in directing spindle assembly towards chromatin. It remains open whether Nup50 already modulates RCC1 activity during this phase of the cell cycle, akin to its newly identified role in nuclear reformation.

## Methods

### Protein expression and purification

*Xenopus laevis* Nup50 full length and fragments were cloned as codon optimized sequences for expression in *E. coli* into a modified pET28a vector with a yeast SUMO solubility tag which is followed by a Tobacco Etch Virus (TEV) cleavage site, a GST-tag or an EGFP-tag. Both mouse Nup50 orthologues were cloned into an pET28a. Proteins were expressed in BL21de3 *E.coli* by auto induction in LB medium at 18°C and purified using Ni-NTA beads (Qiagen). When present, the SUMO-tag was cleaved using TEV protease at 4°C overnight. SUMO-tag and protease were removed using Ni-NTA beads (Qiagen) and the protein of interest dialyzed in a sucrose buffer (250 mM sucrose, 50 mM KCl, 10 mM Hepes-KOH, 2.5 mM MgCl_2_). Ran wild type and the Q69L mutant as well as RCC1 were purified as described [44].

### Antibodies

A Nup50 antibody from Abcam (ab137092) was used at a 1:500 dilution for immunofluorescence in human cells and 1:1000 dilution for western blots, a Nup50 antibody from Abcam (ab85915) at a 1:500 dilution for immunofluorescence in mouse cells and at a 1:1000 dilution for western blot. A MEL28/ELYS antibody from Bio Matrix Research (BMR 00513) was used at a 1:500 dilution for immunofluorescence in human cells and 1:1000 dilution for western blots. The MEL28/ELYS antibody used on mouse cells was a gift from Thomas Schwartz. Mouse monoclonal antibodies mAB414 were from Covance (MMS-120R, used 1:2000 for immunofluorescence and 1:10 000 for detection of Nup62, Nup214 and Nup153 in western blot), His6 antibodies from Roche (11922416001, used 1:1000 for western blotting), GFP antibodies from Roche (11814460001, 1:2000), tubulin from Sigma (T61999, used at 1:10 000 for western blots on human cells), actin from MP Biomedicals (691001, used at 1:1000 for western blots on mouse cells) and ANP32A from Cell Signaling (#15491, D7Z5U used 1:1000, for western blotting). Rabbit polyclonal Nup50 antibodies were generated in rabbits using full-length Xenopus laevis Nup50. Antibodies against Xenopus Nup53 [61], Nup62 [62], MEL28/ELYS [5], Nup153, Importin β and lamin B [19] as well as Ran, RCC1 and importin α [44] were previously described. Antibodies were employed in a dilution of 1:1000 for western blotting and 1:100 for immunofluorescence. Beads for immunodepletion were generated as described [62].

### Pull-down experiments

For pulldown assays, 0.5 µM GST bait proteins (Nup50 full-length or fragments, Nup53 aa 162-267 served as control [63]) were incubated with in a 400 µl of egg extracts for 1h at 4°C, supplemented with 60 µl of 50% slurry of GSH-Sepharose 4B (GE Healthcare) and incubated for another 2h. The sepharose was washed five times with PBS, and bead bound proteins eluted in 30 µl total volume by TEV protease cleavage (0.5 mg/ml) for 1 hour at 25°C. The TEV protease cleaves the GST-fusions between the GST moiety and the bait proteins. The input and elutions were analyzed by western blotting using the indicated antibodies. For quantitation, ECL signals were determined using FiJi, an open-source platform for biological-image analysis [64], background signal were subtracted and the signal ratio of eluate/input was calculated.

Nup50 fragments aa 1-120, 29-120 and 48-120 of human Nup50 were cloned into pCMV-3Tag-1B vector. Both mouse Nup50 paralogues were cloned into pEGFP-N3 and pEGFP-C3 vectors. Xenopus laevis Nup50 and point mutants were cloned into pEGFP-C3 vector. HEK293T cells were cultivated in DMEM medium with 10% fetal bovine serum and 1% penicillin/streptomycin. For FLAG-tag-IPs, pCMV-3Tag-1B vectors including an empty control vector were transfected into the cells by using JetPRIME transfection reagent (PolyPlus Transfection, Illkirch, France) according to the manufacturer indication. After 24 h transfection, the cells were lysed with ice-cold NP-40 lysis buffer (20 mM HEPES-NaOH, pH 7.4, 150 mM NaCl, 0.5% NP-40, 2 mM EDTA) containing protease inhibitor. The cell lysates were incubated with Anti-FLAG M2 Magnetic Beads (Sigma, Taufkirchen, Germany) end-over-end rocking for 2 hours at 4°C. After washing, the beads were used for mass spectrometry-based proteomic analysis or western blotting. Similarly, pEGFP-N3 and pEGFP-C3 based vectors were transfected into HEK293T cells by using JetPRIME transfection reagent. 24 hours late, the cells were lysed with ice-cold NP-40 lysis buffer (see above) and cell lysates were incubated with GFP-Trap Magnetic Beads (ChromoTek, Planegg-Martinsried, Germany) end-over-end rocking for 2 hours at 4°C. After washing, proteins were analyzed by SDS-PAGE and immunoblotting.

### Mass spectrometry

For FLAG-Nup50 aa1-120 interactome analysis, Co-IP experiments (n = 3) were prepared as described above, except that after the immunoprecipitation the beads were washed three times with NP-40 lysis buffer and subsequently three times with lysis buffer w/o detergent. The dried beads from the individual IPs were then processed for MS-analysis as described previously [65]. Briefly, the beads were digested for 1 hour with 5 μg/ml Trypsin (in 2 M Urea, 50 mM Tris-HCl pH 7.5) at RT. The supernatant was then collected in a fresh tube and the beads were washed twice with 2 M urea, 50 mM Tris-HCl pH 7.5 and 1 mM DTT. All supernatants were combined and left to digest o/n at room temperature. The digested peptides were treated with iodoacetamide, acidified, desalted (homemade C18-tips) and lyophilized and stored at −80°C. Prior to mass spectrometry analysis, the peptides were resuspended in 3% formic acid (FA)/5% acetonitrile (ACN) and loaded onto a nanoLC system (RSLCnano, Thermo Scientific). First, the Peptides were trapped for 10 min on a precolumn (Acclaim PepMap100, C18, 5 μm, 100 Å, 300 μm i.d. × 5 mm, Thermo Scientific) and subsequently separated using an analytical column (Easyspray 50 cm column (ES803) at 45°C; Thermo Scientific) employing a 125 min gradient (0–10 min: 5% buffer B (buffer A: 0.1 % FA; buffer B: 80 % acetonitrile, 0.1 % FA), 10–60 min: 5–20 % buffer B, 60-98 min: 20-35 % buffer B, 98-101 min: 35-99 % buffer B, 101-106 min: 95 % buffer B, 106-109 min: 95-5 % buffer B, 109-125 min: 5 % buffer B) with a spray voltage of 2 kV and the capillary temperature at 250 °C. All samples were analyzed on a Q Exactive plus mass spectrometer (Thermo Scientific) in data dependent mode. Full MS settings were: 70,000 resolution; AGC target, 1e6; maximum injection time, 50 milliseconds; scan range, 350-1600 m/z. dd-MS2 settings were: 17,500 resolution; AGC target: 1e5; maximum injection time: 55 milliseconds; top 20 precursor fragmentation; isolation window, 2.0 m/z; collision energy, 27. dd settings were: minimum AGC, 5e2; 20 second dynamic exclusion; only 2+ to 5+ peptides were allowed.

Analysis of the raw data was done using MaxQuant (version 1.6.17.0) with the built-in Andromeda search engine [66]. The search was conducted against the human SwissProt database version 12/2020 (only reviewed and canonical sequences) and MaxQuant default settings (including the mass tolerance) were used. Specific settings: Trypsin as the specific protease (two missed cleavages); Carbamidomethylation (Cys) as fixed modification; Oxidation (Met) and N-terminal protein acetylation as variable modifications. The false discovery rate was 0.01 on both peptide and protein level and the minimum peptide length was seven amino acids. Quantification was done using the label free quantitation algorithm from MaxQuant.

The proteinGroups.txt result file (Sup. Tab. 1) from the MaxQuant search was then subjected to analysis using the Perseus software suite (version 1.6.14.0) [67]. The LFQ intensities from all biological replicates were used as the main columns. Filtering of the protein list was carried out for reversed hits, contaminants, and “only identified by site” entries. Further requirements for protein inclusion included minimum 1 unique peptide and 2 total peptides (razor + unique). The 3 individual biological replicates were grouped as NUP and ctrl and the data was then log2-transformed. Proteins were only included in the final data set, if they were identified in all replicates in at least one group (min 3 in one group). The remaining list was subjected to imputation of missing values based on normal distribution (Perseus default settings). The data was further analyzed using the two-sample tests option. The resulting file was used for generation of the Volcano plots (Fig. 8A). The settings for the volcano plot were: p-value<0.01 and a ratio of >4-fold change (>2 (−log (10) p-value) and >2 (log (2) ratio).

### *In vitro* nuclear assembly

Preparation of high speed interphase extracts, sperm heads, floated labelled and unlabeled membranes required for *in vitro* nuclear assembly as well as immunofluorescence experiments were carried out as described in [28], nuclear transport assays as in [62]. Fluorescence images were acquired using a confocal microscope [FV1000; Olympus; equipped with a photomultiplier (model R7862; Hamamatsu)] using 405-, 488- and 559-nm laser lines and a 60× NA 1.35 immersion oil objective lens or a Zeiss LSM710 confocal microscope equipped with a Plan-Apochromat 63×/1.4 immersion oil objective and 405 nm, 488 nm and 561 nm lasers, using ZEN software. Annulate lamellae assays were performed in 45 μl of Xenopus egg extract, 5μl of floated membrane and an energy regenerating system containing [28]. Protein of interest were added at 10 μM final concentration in a reaction volume of 60 μl and incubated for 4 h at 20°C. Processing of the samples were carried out as previously described [19]. Reactions were stopped by adding 1 ml of sucrose buffer (250 mM sucrose, 50 mM KCl, 10 mM Hepes-KOH, 2.5 mM MgCl_2_), spun for 10 min at 7500 g. The pellet was re-suspended in 10 μl SDS-sample buffer and analyzed by SDS-PAGE and Western blotting using mAB414 antibodies at 1:10 000 dilution. ECL signals were quantified using FiJi, measuring the mean intensity of the Nup62 signal in each condition, subtracting the background signal and normalizing to the control.

### Cellular experiments

HeLa, HeLa stably expressing H2B-mCherry, HEK293T and NIH-3T3 cells were grown in Dulbecco’s Modified Eagle’s Medium (DMEM, Gibco) with pyruvate supplemented with 10% of decomplemented fetal bovine serum (Gibco) and 100U/ml of penicillin (Gibco) and 100μg/ml of streptomycin (Gibco). Cells were incubated at 37 °C in a 5% CO_2_ atmosphere. siRNA against Nup50 (SI00663236) was obtained from Qiagen. The siRNA against mouse Nup50 (156930) was purchased from Ambion Life Technologies. The siRNA against the mouse Nup50B was custom made with the following sequence: 5’ GAAGCCAGCAUCUGCCAAAtt 3’ and purchased from ambion life technologies. MEL28/ELYS siRNA was obtained from Sigma Aldrich (3976251-F). Cell transfection was performed in 24 well plate with Lipofectamin RNAimax from Thermofisher scientific. Lipofectamine and siRNA were mixed in OptiMEM (Gibco), incubated at least 20 min at room temperature and put in a well. 520 μl of cell suspension in growing medium was added on the top of 80 μl of siRNA solution to obtain a final concentration of 20 nM of siRNA and 20 000 cells per well. Cells were incubated 72 hrs at 37 °C in a 5% CO_2_ atmosphere.

For localizing different EGFP-Nup50 fragments and mutants in cells, HeLa cells at a confluence of 20-30% were transfected with the respective EGFP-C3 constructs using FuGENE6 transfection reagent (Roche) following the manufacturer’s instructions. After 36h, the cells were washed three times with PBS and for 1sec preextracted using PBS supplemented with 0.1% (v/v) Triton-X100 (Merck) to reduce the nucleoplasmic EGFP signals. After fixation with 4% paraformaldehyde in PBS for 20 minutes, the chromatin was stained with RedDot™2 Far-Red Nuclear Stain (Biotium, 1:100 diluted in PBS), and the samples mounted using Vectashield Mounting Medium and analyzed on an FV1000; Olympus confocal microscopy (see above).

The mAB414 signal quantified using FiJi. Total nuclear areas were automatically selected using the threshold option and inner nuclear area were manually drawn and subtracted to obtain the nuclear rim area and calculated its mean signal intensity. For each experiment the mAB414 signal was divided by the average signal of the control condition, resulting in a distribution of signal intensity centered on 1 in the control condition. Signal distributions were then plotted on a box plot and whisker graph showing the median, interquartile distribution, minimum and maximum values and a Mann-Whitney statistical test was performed using GrapPad.

### Life cell imaging

HeLa cells stably expressing H2B-mCherry generated as in [68] and cultured as above were seeded in 8 well μ-slide chambers (Ibidi) and next day transfected with 9 ng/μl from the indicated eGFP-fusions (pEGFP-C3 constructs expressing wild type and mutant mouse Nup50 proteins) using jetPRIME (Polyplus, Illkirch, France). The cells were imaged 24 hours post transfection with a spinning disk confocal microscope Ti2 Eclipse (Nikon) equipped with a LED light engine SpectraX (Lumecor, Beaverton, Oregon, USA) and GFP/mCherry filter sets, a Lambda Oil 60× NA 1.3 objective and environmental control system UNO-T-H-CO2 (Okölab, Ottaviano, NA, Italy). AR-Elements software (Nikon) equipped with a software based autofocus module was used to perform confocal fluorescence imaging of the single best-in-focus optical section of cells in metaphase every three minutes during approximately 2 hours. Image galleries from cell trajectories during mitotic exit were assembled using FiJi and mounted for figures using Inkscape (Free Software Foundation, Inc. Boston, USA).

### DNA-beads experiments

The MCP1 plasmid [38] was digested by BamHI and NotI and the digestion product was purified. 30 μg of DNA was biotinylated using 0.36 U of Klenow fragment, 63μM of biotin-18-sUTP, 63μM of biotin-14-dATP, 90 μM of thio-CTP and 90 μM of thio-GTP. The DNA was purified in a CHROMA SPIN column according to the manufacturer’s instructions. The biotinylated DNA was coupled to Dynabeads Kilobinders (Thermofisher Scientific) following the manufacturer’s instructions. DNA-coupled magnetic beads were incubated with protein in a table top shaker 3 hours at 20°C at 350 rpm. After washing, beads were collected in 10 μl of egg extract or sucrose buffer (250 mM sucrose, 50 mM KCl, 10 mM Hepes-KOH, 2.5 mM MgCl_2_) and incubated with DAPI (Roth) at 1μg/ml 10min at room temperature. 3 μl were sampled on a 3 well Diagnostika slides (X1XER303B) from Thermo scientific for observation on an Zeiss LSM710 confocal microscope equipped with a Plan-Apochromat 63×/1.4 Oil objective and 405 nm and 488 nm lasers, using ZEN software..

### RCC1 guanine nucleotide exchange assay

GEF assays were adapted from [69]. Recombinant Ran was loaded with MANT-GDP (Jena Bioscence, NU-204) in a low magnesium loading buffer (20 mM HEPES-NaOH, 50 mM NaCl, 0.5 mM MgCl_2_, 10 mM EDTA, 2 mM DTT and 20 fold molar excess of MANT-GDP over Ran). Exchange for GppNHp was performed. Ran MANT-GDP to GppNHp (Jena Bioscence, NU-401) exchange experiments were carried out with 2 μM of Ran in a volume of 50 μl in a black 96 well glass bottom plate in a nucleotide exchange buffer (40 mM HEPES-NaOH, 50 mM NaCl, 10 mM MgCl_2_ and 2 mM DTT) in the presence of 2mM GppNHp excess. The assays were performed with an excitation wavelength of 360 nm and emission at 440 nm in a SpectraMax iD3 spectrophotometer from Molecular Devices every 15 seconds.

### Sequence alignment

Human (ENST00000347635.9), *Xenopus laevis* (BC077201.1), *Xenopus tropicalis* (NM_001016628.2), mouse (A: ENSMUST00000165443.4; B: ENSMUST00000090061.6), rat (A: ENSRNOT00000018155.6; B: ENSRNOG00000024560), chicken (ENSGALT00000022998.6), zebrafish (ENSDART00000004739.8), fruitfly (FBtr0088783), red flour beetle (TC007383_001), *S. cerevisiae* (YLR335W), *S. pombe* (SPCC18B5.07c.1) and *S. japonicus* (EEB09106) transcript sequences were retrieved from Ensembl or Genebank and aligned by ClustalO with the Seaview software [70]. Sequence identity scores were calculated using the sequence identity and similarity (SIAS) webpage of the computational university of Madrid with the default settings.

## Author Contributions

Conceptualization, G.H, U.K. and W.A.; formal analysis, G.M., P.D.M., C.G., R.S., A.S., D.M.-A. C.P., M.L.; investigation, G.M., P.D.M., C.G., R.S., A.S., D.M.-A. C.P., D.M.-A., H.L.., G.H., A.S., H.L., A.K.S. A.M., M.T. and M.W.; resources, B.L. and H.L.; writing—original draft preparation, G.H. and W.A.; writing—review and editing, G.H. and W.A.; visualization, G.H. D.M.-A. and C.P.; supervision, D.M.-A.,U.K. and W.A.; project administration, W.A.; funding acquisition, U.K. and W.A. All authors have read and agreed to the published version of the manuscript.

## Acknowledgments

Mass Spectrometry was done at the Proteomic Facility and Confocal images were acquired at the Confocal Microscopy Facility, core facilities of the Interdisciplinary Center for Clinical Research (IZKF) Aachen within the Faculty of Medicine at RWTH Aachen University. This work was supported by grants from the German Research Foundation (to W.A. AN377/7-1), the Swiss National Science Foundation (to UK 310030_184801)

