## Supplemental figures for "The nucleoporin Nup50 activates the Ran guanyl-nucleotide exchange factor RCC1 to promote mitotic NPC assembly"

**Figure S1: Characterization of Nup50 antisera, recombinant protein and depletion phenotype**

(A) Western blot against Xenopus Nup50 on Mock treated (control, left lane) or Nup50 depleted (right lane) Xenopus egg extract.

(B) Confocal microscopy images of nuclei assembled for 120 min in mock depleted, Nup50 depleted (ΔNup50), and Nup50 depleted *Xenopus* egg extracts supplemented with recombinant Nup50. Then 5µM recombinant EGFP-M9-M10 as import substrate or EGFP-M9-NES as export substrate [71] were added, the later in the absence or presence of 300 nM leptomycin. After 120 min , nuclei were fixed in 4% PFA and 0.5 % glutaraldehyde, membranes were pre-labelled with DiIC18 (1,1'-Dioctadecyl-3,3,3',3'-Tetramethylindocarbocyanine Perchlorate, red) and chromatin was stained with DAPI (4′,6-Diamidin-2-phenylindol, blue).

(C) Coomassie gel showing recombinant Nup50 purification. Lanes show 1, 2 and 4 µl of Nup50 form left to right.

**Figure S2: Nup50 interacts with MEL28/ELYS**

(A) Western blot analysis of immunoprecipitation of MEL28/ELYS (center panel), Nup50 (right panel) or input (left panel) using mitotic (M) or interphase (I) Xenopus extract using MEL28/ELYS (upper lanes) or Nup50 (lower lanes) antibodies.

(B) Quantitation of (A) the mean signal of the Input, MEL28/ELYS IP and Nup50 IP are plotted the left to the right. The white bar represents the amount of Nup50 protein and the grey bar represents the amount of MEL28/ELYS from n=4 independent experiments. Individual data points are indicated, data normalized the input condition.

(C) Western blot analysis of Mock treated, MEL28/ELYS depleted or Nup50 depleted Xenopus egg extracts with MEL28/ELYS and Nup50 antibodies. The importin α-export factor CAS serves as loading control.

(D) Relative quantification of MEL28/ELYS and Nup50 proteins in depleted extract. Quantitation in mock treated extracts, MEL28/ELYS depleted extracts and Nup50 depleted extracts are respectively plotted from the left to the right. The white bar represents the amount of Nup50 protein and the grey bar represents the amount of MEL28/ELYS from n=7 independent experiments. Individual data points are indicated, data normalized to the mock condition.

**Figure S3: Time course of Nup50 and MEL28/ELYS chromatin recruitment**

(A, B) Demembranated sperm chromatin was preincubated in Xenopus egg extract for 10 min. At time point zero, membranes were added to the reaction. Reactions were stopped at the indicated time points by fixation and analyzed by confocal microscopy after immunofluorescence with α-Nup50 (A), α-MEL28/ELYS (B), and mAb414 (A, B). Chromatin was stained with DAPI (left column in A and B). Scale bar: 10 µm.

**Figure S4: Alignment of Metazoan Nup50 144-189 with Nup2 from different yeast species**

Sequence alignment of Nup50 144-189 (*X. laevis* numbering) with Nup2/Nup61, species are indicated on the left. The color scheme indicates the type of amino acids according to the alignment software default setting.

**Figure S5: Localization of Nup50 mutants in in vitro nuclear assembly assays**

Confocal microscopy images of nuclei assembled for 120 min in mock depleted, Nup50 depleted (ΔNup50), and Nup50 depleted *Xenopus* egg extracts supplemented with recombinant wild type Nup50 or different mutants. Nuclei were fixed in 4% PFA and 0.5 % glutaraldehyde, stained for NPCs (mAB414), Nup50 and the chromatin (DAPI). Scale bar: 10 µm.

**Figure S6: Nup50 binds membranes**

(A) Recombinant EGFP-tagged Nup50 (green) was incubated with giant unilamellar vesicles (GUVs), which were labeled with the lipophilic dye DiIC18 (1,1'-Dioctadecyl-3,3,3',3'-Tetramethylindocarbocyanine Perchlorate, red). Confocal images of individual channels and the overlay is shown. Scale bar: 10 µm.

(B) Binding assay of recombinant full-length Nup50, a Nup98 fragment (676-863, negative control) and a Nup153 fragment (aa 1-149, positive control) on small 30 nm, 100nm or 400nm liposomes. The percentage of floated protein is shown. Bars show the average of three independent experiments, individual data point are indicated.

(C) Liposome floatation assay using 100 nm liposomes of recombinant full-length Nup50 or different Nup50 fragments. The percentage of floated protein is shown. Bars show the average of at least three independent experiments, individual data point are indicated.

**Figure S7: Nup50 N-terminus interacts with RCC1**

(A) HEK293T cells were transfected with empty control vector, different FLAG-Nup50 N-terminal fragments aa 1-120 and aa 48-120 (of the human sequence, respectively corresponding to the 1-114 or 48-114 residues of the Xenopus sequence.) or different double mutants designed to compromise RCC1 binding. 24h post transfection cells were lysed, FLAG-tagged proteins immunoisolated and analyzed by western blotting, with 10% of the inputs loaded.

(B) Xenopus egg extract were supplemented with 5 µM RanQ69L, RCC1, Nup50 wild type and fragments as well indicated combinations of 5 µM Nup50 and 5 µM RCC1. After 90 min, annulate lamellae were isolated by centrifugation and quantified by western blotting with mAB414 antibody. Quantitation shows the relative Nup62 signal as a mean from at least three experiments, normalized to the buffer control. Individual data points are indicated.

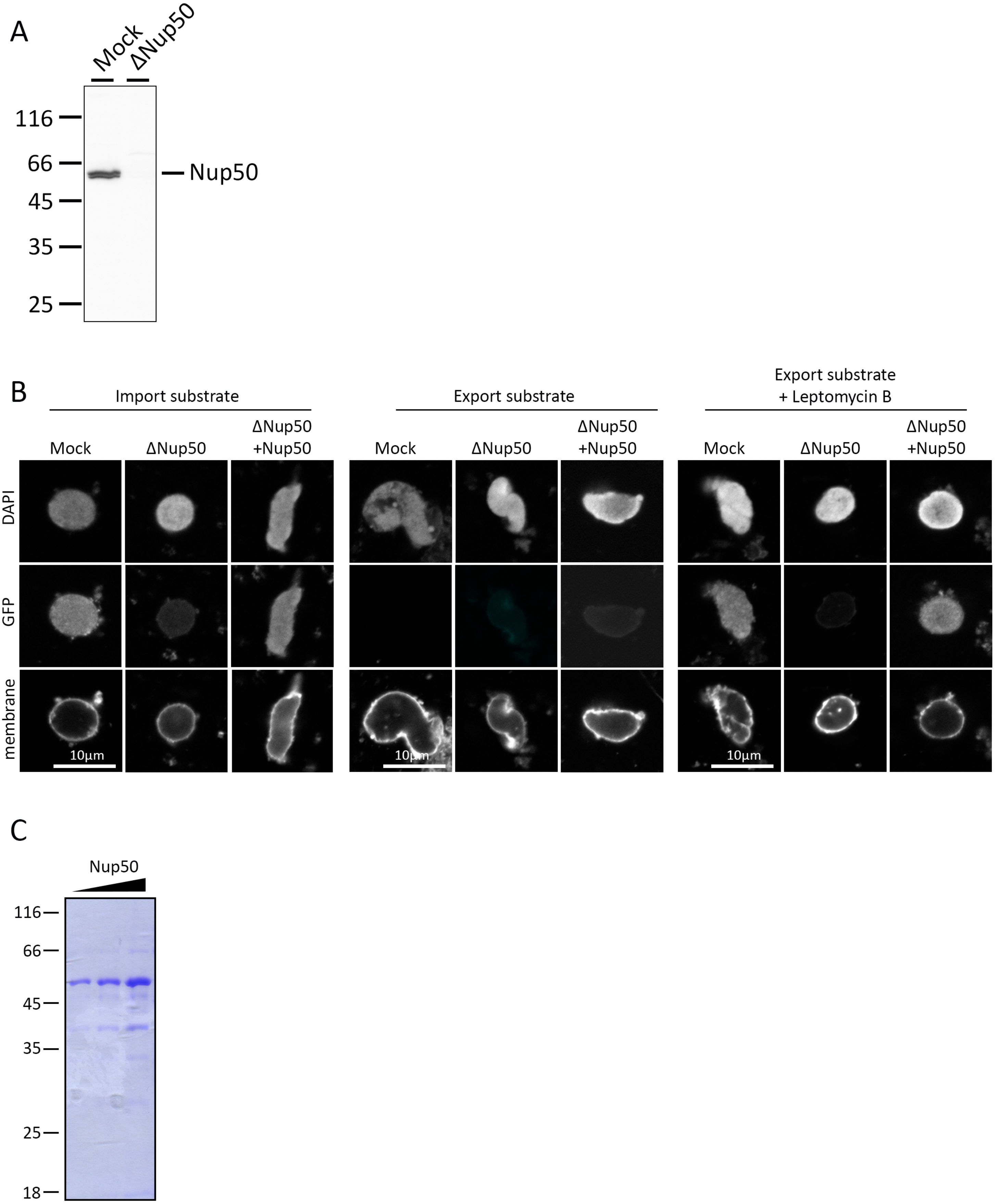
A
Mock
ΔNup50
116
66
45
35
25
Nup50
B
Import substrate
Export substrate
Export substrate
+ Leptomycin B
Mock
ΔNup50
ΔNup50
+Nup50
DAPI
GFP
membrane
10µm
C
Nup50
116
66
45
35
25
18

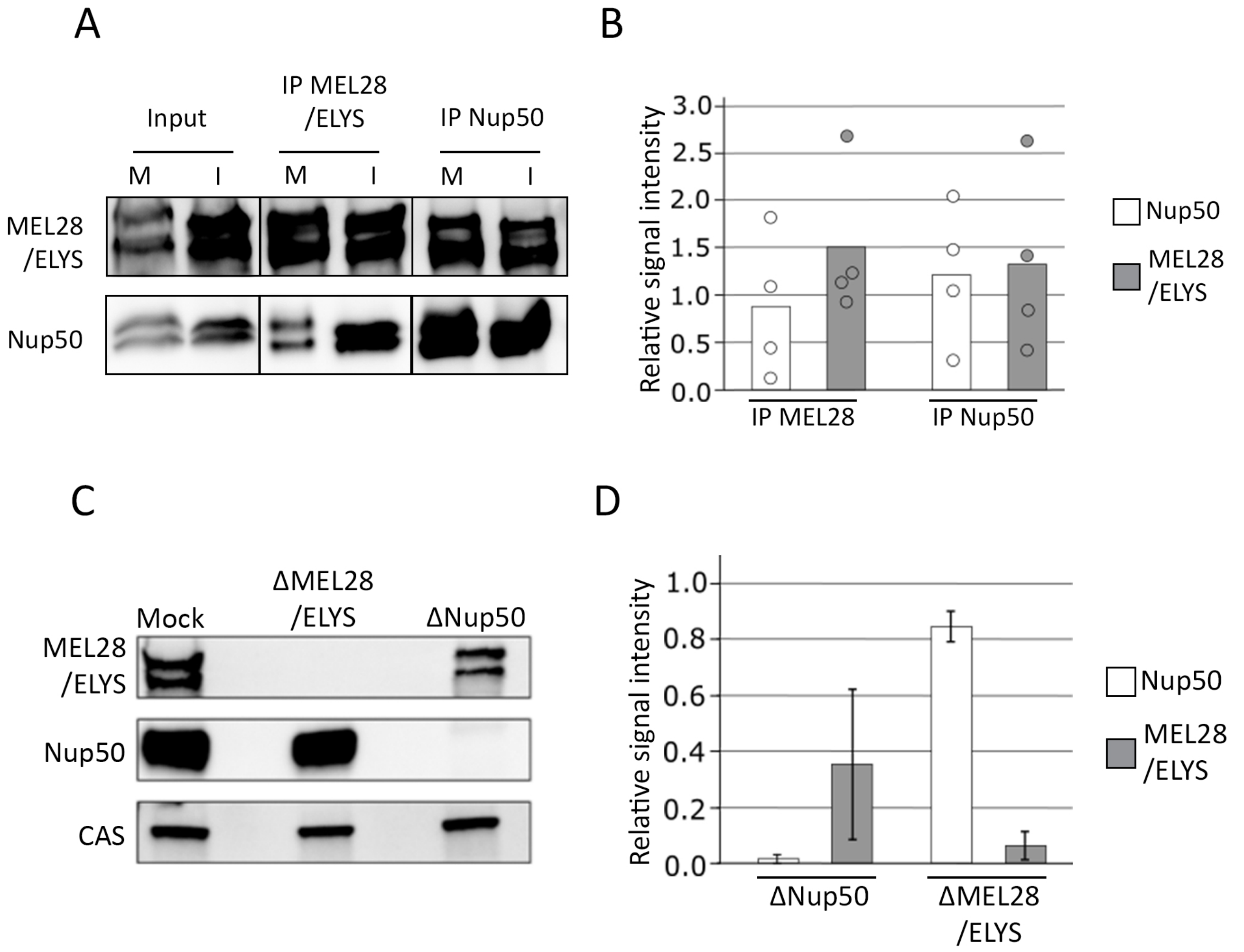
A
Input
IP MEL28 /ELYS
IP Nup50
M
I
M
I
M
I
MEL28 /ELYS
Nup50
B
Relative signal intensity
3.0
2.5
2.0
1.5
1.0
0.5
0.0
IP MEL28
IP Nup50
Nup50
MEL28 /ELYS
C
Mock
ΔMEL28 /ELYS
ΔNup50
MEL28 /ELYS
Nup50
CAS
D
Relative signal intensity
1.0
0.8
0.6
0.4
0.2
0.0
ΔNup50
ΔMEL28 /ELYS
Nup50
MEL28 /ELYS

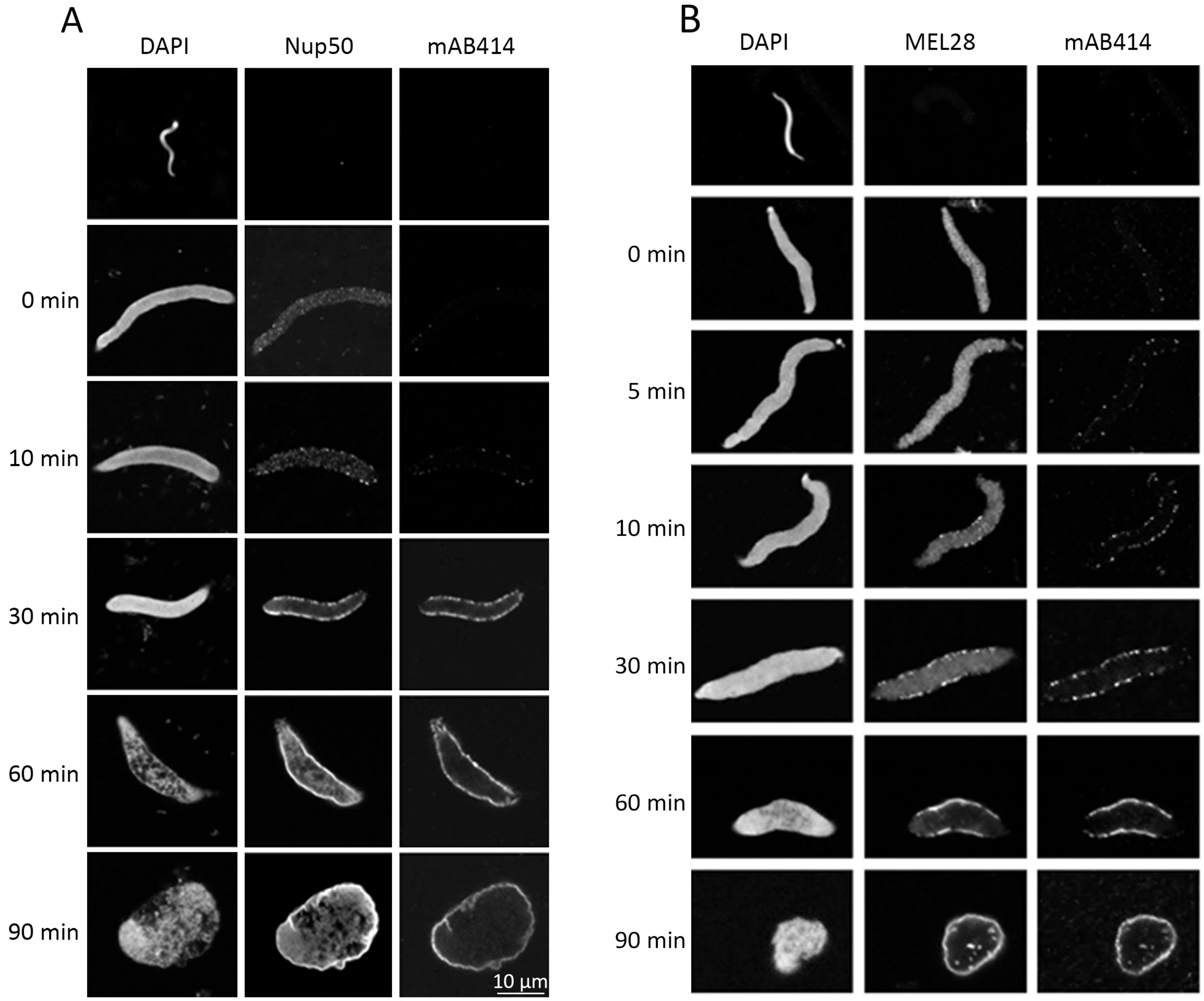
A
DAPI
Nup50
mAB414
0 min
10 min
30 min
60 min
90 min
10 µm
B
DAPI
MEL28
mAB414
0 min
5 min
10 min
30 min
60 min
90 min

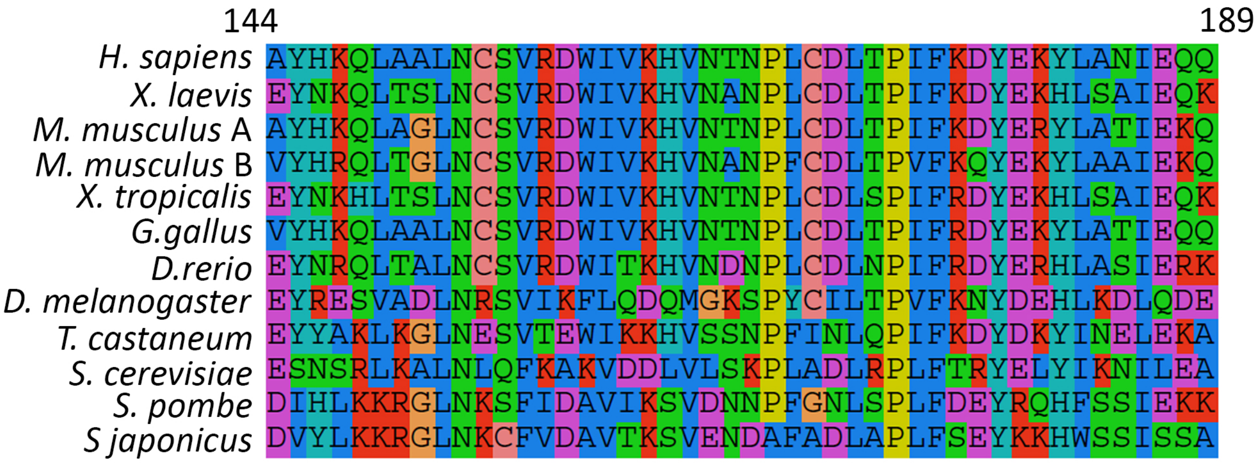
144
189
H. sapiens AYHKQLAALNCSVRDWIVKHVNTNPLCDLTPIFKDYEKYLANIEQQ
X. laevis EYNKQLTSLNCSVRDWIVKHVNANPLCDLTPIFKDYEKHLSAIEQK
M. musculus A AYHKQLAGLNCSVRDWIVKHVNTNPLCDLTPIFKDYERYLATIEKQ
M. musculus B VYHRQLTGLNCSVRDWIVKHVNANPFCDLTPVFKQYEKYLAAIEKQ
X. tropicalis EYNKHLTSLNCSVRDWIVKHVNTNPLCDLSPIFRDYEKHLSAIEQK
G.gallus VYHKQLAALNCSVRDWIVKHVNTNPLCDLTPIFRDYEKYLATIEQQ
D.rerio EYNRQLTALNCSVRDWITKHVNDNPLCDLNPIFRDYERHLASIERK
D. melanogaster EYRESVADLNRSVIKFLQDQMGKSPYCILTPVFKNYDEHLKDLQDE
T. castaneum EYYAKLKGLNESVTEWIKKHVSSNPFINLQPIFKDYDKYINELEKA
S. cerevisiae ESNSRLKALNLQFKAKVDDLVLSKPLADLRPLFTRYELYIKNILEA
S. pombe DIHLKKRGLNKSFIDAVIKSVDNNPFGNLSPLFDEYRQHFSSIEKK
S japonicus DVYLKKRGLNKCFVDAVTKSVENDAFADLAPLFSEYKKHWSSISSA

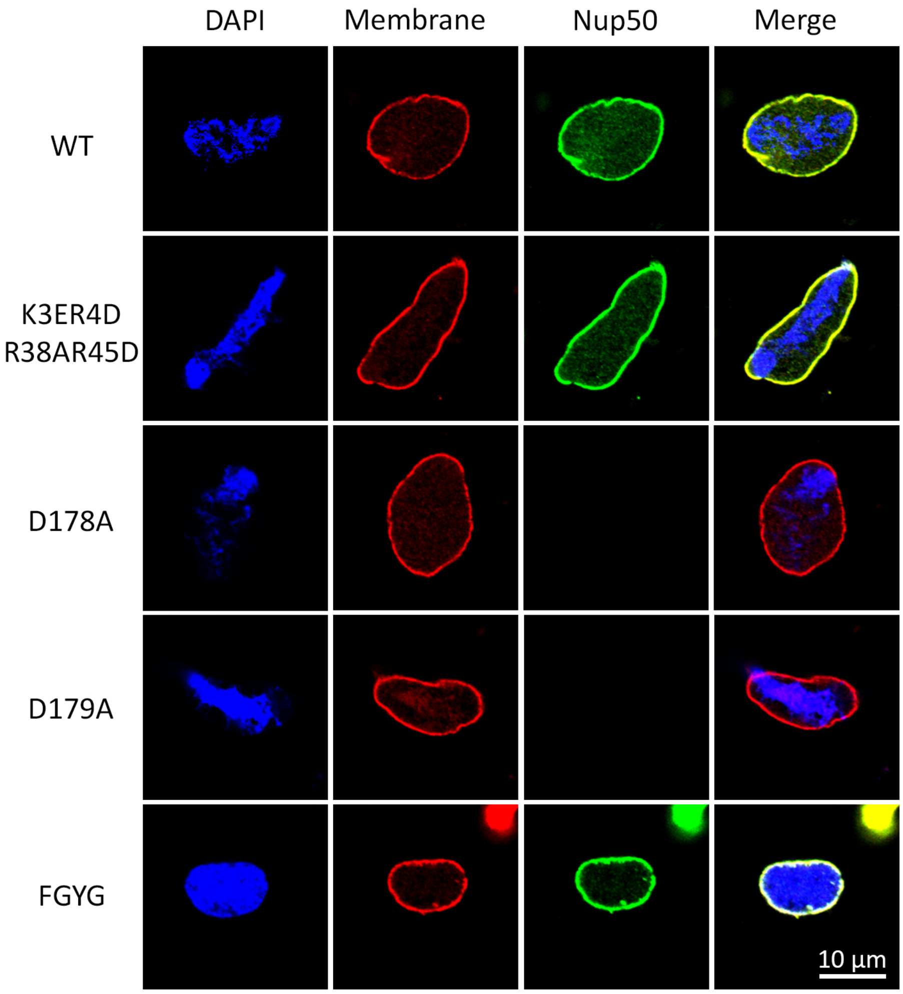
DAPI
Membrane
Nup50
Merge
WT
K3ER4D
R38AR45D
D178A
D179A
FGYG
10 µm

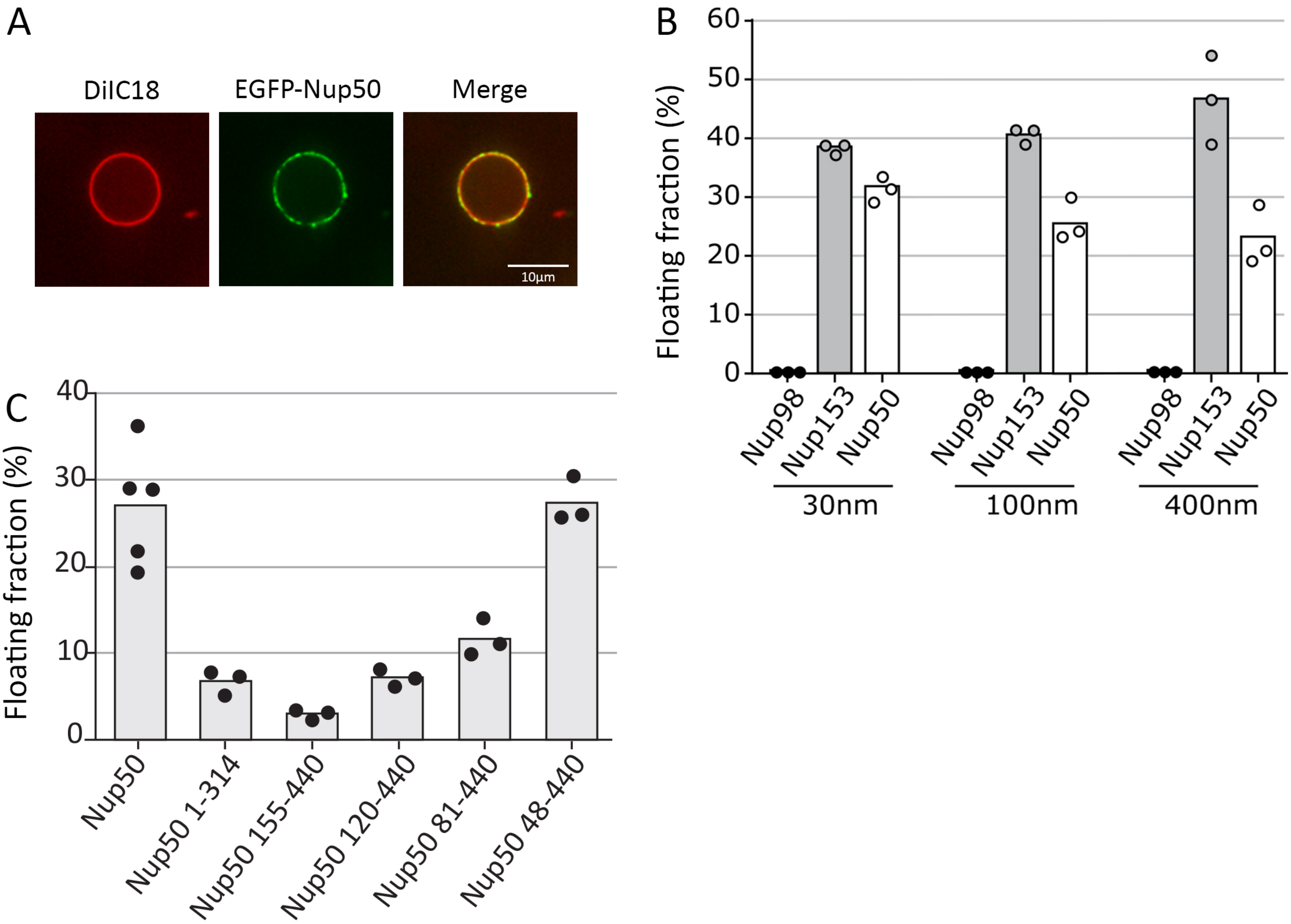
A
DiIC18
EGFP-Nup50
Merge
10µm
B
Floating fraction (%)
60
50
40
30
20
10
0
Nup98
Nup153
Nup50
30nm
100nm
400nm
C
Floating fraction (%)
40
30
20
10
0
Nup50
Nup50 1-314
Nup50 155-440
Nup50 120-440
Nup50 81-440
Nup50 48-440

A

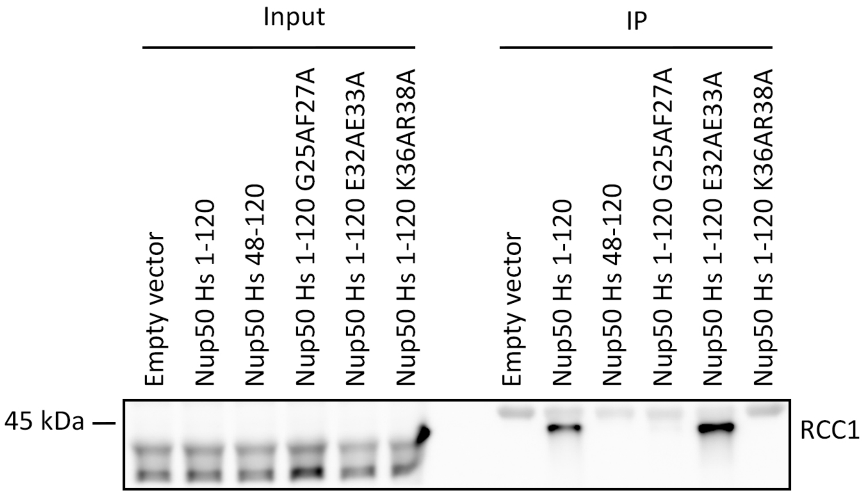
Input
IP
Empty vector
Nup50 Hs 1-120
Nup50 Hs 48-120
Nup50 Hs 1-120 G25AF27A
Nup50 Hs 1-120 E32AE33A
Nup50 Hs 1-120 K36AR38A
Empty vector
Nup50 Hs 1-120
Nup50 Hs 48-120
Nup50 Hs 1-120 G25AF27A
Nup50 Hs 1-120 E32AE33A
Nup50 Hs 1-120 K36AR38A
45 kDa
RCC1

B

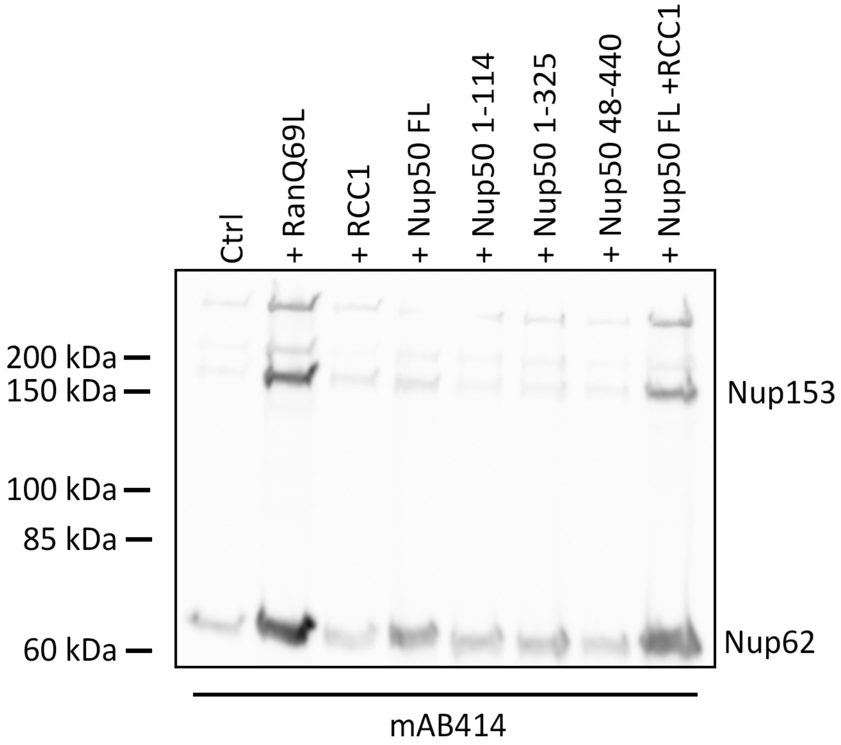
Ctrl
+ RanQ69L
+ RCC1
+ Nup50 FL
+ Nup50 1-114
+ Nup50 1-325
+ Nup50 48-440
+ Nup50 FL +RCC1
200 kDa
150 kDa
100 kDa
85 kDa
60 kDa
Nup153
Nup62
mAB414

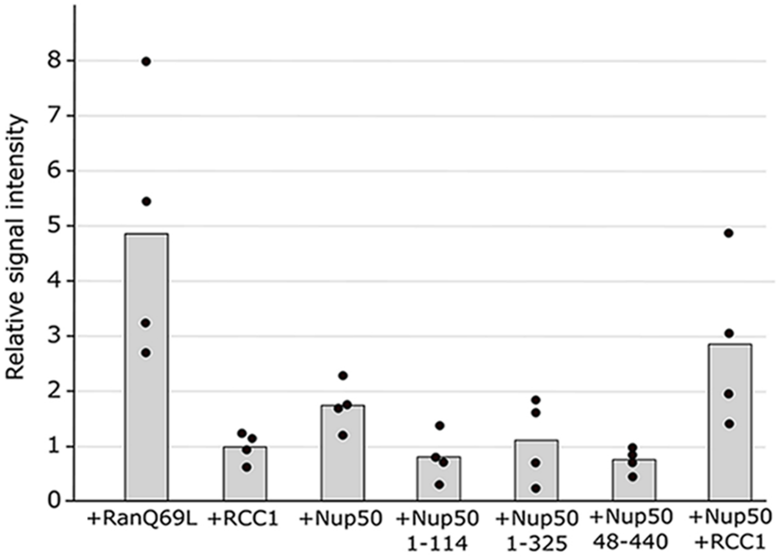
Relative signal intensity
0
1
2
3
4
5
6
7
8
+RanQ69L
+RCC1
+Nup50
+Nup50 1-114
+Nup50 1-325
+Nup50 48-440
+Nup50 +RCC1
